# Navigating the Landscape of Fear

**DOI:** 10.1101/2024.08.15.608007

**Authors:** Marissa Gee, Nicolas Gonzalez-Granda, Sunay Joshi, Nagaprasad Rudrapatna, Anne Somalwar, Stephen P. Ellner, Alexander Vladimirsky

**Affiliations:** Center for Applied Mathematics, Cornell University, Ithaca, NY; Division of Biostatistics and Health Data Science, School of Public Health, University of Minnesota, Minneapolis, MN; Graduate Group in Applied Mathematics and Computational Science, University of Pennsylvania, Philadelphia, PA; Oden Institute for Computational Engineering and Sciences, The University of Texas at Austin, Austin, TX; Department of Ecology and Evolutionary Biology, Cornell University, Ithaca, NY; Department of Mathematics, Cornell University, Ithaca, NY

## Abstract

1. Animals searching for food must navigate complex landscapes with varying terrain, food availability, predator activity, and shelter. Where and when should they gather food? To what extent should they engage in anti-predator behaviors such as vigilance or seeking refuge if a predator is detected? Optimal foraging theory (OFT) posits that animals balance potentially conflicting goals (such as feeding versus escaping predation) by making decisions that maximize some expected utility or reward. However, OFT models have generally considered highly simplified landscapes, either ignoring spatial variability or assuming that the habitat consists of discrete, internally uniform habitat patches. As a result, OFT has largely avoided the question of how animals should move from one potential feeding area to another, or between feeding areas and refuges.
2. We develop methods based on stochastic dynamic programming to find optimal foraging strategies, including optimal movement paths, in a continuous landscape with spatially varying predation risk. Our approach accounts for switching from foraging to escape behavior when pursued by a predator. Because contingent escape paths are considered for all visited locations, they influence the optimal foraging path even before threats are encountered. The optimal strategy thus depends on the animal’s level of hunger, the distribution of food, and the perceived threat distribution. The realized path is further influenced by actual predator encounters.
3. We illustrate our approach with two numerical examples: the first hypothetical with two food-abundant regions accessible only via high-risk areas, the second based on empirical studies on foraging Samango monkeys, *Cercopithecus albogularis schwarzi*. We find that the shape of the forager’s utility function (risk-averse, risk-neutral, or with state-dependent risk sensitivity) affects not only its choices of where to feed, but also the optimal paths to and from each feeding ground.
4. Our methods make it possible to compare properties of observed foraging trajectories with those predicted for different goal functions. Foraging trajectories can then provide additional information, along with other behavioral choices, about what quantity, if any, animals aim to optimize while foraging.

## Introduction

Foraging animals make decisions every day that impact their chances of survival and reproductive success. Where, when, and for how long should they gather food? To what extent should they engage in anti-predator behaviors such as vigilance or seeking refuge if a predator is detected? These decisions depend on balancing the animal’s short-term and long-term goals, and on the information available about the food abundance and predator density at different locations. Optimal Foraging Theory (OFT), [Stephens and Krebs, 1987] seeks to understand these decisions by viewing animals as rational decision makers, and interpreting observed choices of diet and foraging locations as the best among the available options for maximizing the organism’s fitness (transmission of genes to subsequent generations). Understanding the basis for foraging decisions can shed light on how animals evaluate and trade off between conflicting goals when foraging, inform the design of experiments to test inferences about the basis for foraging decisions, and provide predictions of how behavior may change as a result of a changing environment [Reimer et al., 2019].

Early work in OFT assumed that a forager seeks to maximize its net rate of energy gain [Stephens and Krebs, 1987]. These models examined questions such as how a forager should decide which food items to eat (or attempt to eat) if they are encountered, and how it should exploit different food patches in a patchy environment (see [MacArthur and Pianka, 1966, Emlen, 1966, Charnov, 1976]). While early rate-maximization models focused entirely on energy gain, models of risk-sensitive foraging (e.g., [Caraco, 1980, Stephens, 1981, Houston and McNamara, 1982, Lima and Dill, 1990, Werner and Gilliam, 1984, Gilliam and Fraser, 1987]) were able to incorporate the mortality risk associated with foraging decisions. However, rate-maximization theories cannot reflect how an animal’s priorities might change with its *state* (e.g., its physical location, or how close it is to starvation).

To address these limitations, [Mangel and Clark, 1986, McNamara and Houston, 1986] proposed models that explicitly track the forager’s state, and include randomness in both food acquisition and possible death through predation. These models assume that a forager seeks to maximize the expected reward accrued during a modeled time period, where the reward encodes reproductive success during the time period, plus expected future success based on the forager’s state at the final time. State-dependent optimal decisions are found numerically through stochastic dynamic programming (SDP), versions of which were later used to analyze many theoretical questions about animal behavior [Newman, 1991, Houston et al., 1993]. The SDP framework has seen numerous empirical applications including birds deciding whether to flock or find a mate [Szekely et al., 1991], tuna foraging near ocean fronts [Kirby et al., 2000], and polar bears preparing for hibernation and reproduction [Reimer et al., 2019]. However, in almost all cases the forager’s current energy reserve level is the only state variable. The choice of the next food item or feeding area does not take into account the animal’s current location, or what path it will take to its chosen target.

In this paper, we take a first step in generalizing the state-based dynamic programming approach of classical optimal foraging theory in order to characterize optimal paths for an animal foraging in a spatially continuous habitat. That is, we try to characterize how a forager should navigate through a *landscape of fear*. The term “landscape of fear” has been used in multiple ways, but it generally refers to the spatial distribution of predation risk, real or perceived, throughout a habitat (e.g., [Brown, 1988, Brown et al., 1999, Pinti et al., 2022, Laundré et al., 2001, Gaynor et al., 2019]). Studies have examined how the introduction of predators affects the spatial distribution of prey animals [Laundré et al., 2001, Raynor et al., 2021], spatio-temporal distributions of prey [Kohl et al., 2018], and the relative impact of predation and food availability on utilization of the environment [Coleman and Hill, 2014]. Optimal foraging theory is potentially well-posed to contribute to the analysis of these data, but must first overcome the challenges associated with finding optimal decisions about movement in a continuous habitat where food availability and predation risk are both spatially varying.

Some previous optimal foraging models sought to incorporate spatial dynamics, though with significant simplifications. [Rands, 2017] considers a forager collecting food and avoiding a predator in a discretized 1-dimensional habitat. Interactions between the forager and a predator are modeled explicitly, and the forager must decide not only when to feed and when to seek refuge, but also where to position itself when it is not optimal to do either of those things. [Fiksen et al., 1995] proposed a spatially explicit optimal foraging model of capelin in the Barents sea. The authors estimate food availability, temperature, and mortality due to predation for a complex environment to predict how the distribution of capelin will evolve in time. However, they focus on a timescale of months to years, and thus do not attempt to recover detailed capelin trajectories nor capture any short-term changes in their behavior induced by predator encounters. In contrast, we seek to solve a spatially explicit optimization problem in a continuous habitat with enough temporal resolution to characterize specific forager paths, and abrupt changes to those paths when they are spotted by a predator.

In all SDP-style ecological models, one of the vexing questions is the appropriate optimization criterion. One critique of early optimal foraging models was that the maximization of energy rate gain is not always a reasonable proxy for reproductive success [Stephens, 1981]. Previous work has shown that the shape of the utility/reward function can impact foraging behavior [Newman, 1991]. Here, we do not specify a single reward function but instead contrast foraging behaviors for several different functional forms of the reward.

In the following section, we propose a model of a forager’s movement and energy dynamics based on differential equations, and a stochastic model of the impact of predators. We assume that the forager seeks to maximize an expected terminal utility and present three possible functional forms that encode different risk preferences. Treating this as an optimal control problem, we derive a set of Hamilton-Jacobi-Bellman partial differential equations that we solve numerically to find the forager’s optimal policy. We use our model to analyze two examples: one mimicking the risk/reward trade-offs in an environment with two main feeding grounds, the other nspired by an empirical study of foraging Samango monkeys in the Soutpansberg Mountains in South Africa [Coleman and Hill, 2014]. Quantitative predictions about foraging animals would require more information than is typically available, and in many cases a more detailed model than we present here. Nonetheless, our model and methods can provide qualitative insight into what behavior would arise from optimizing different utility functions. This in turn can inform the design of experiments or observational studies that seek to infer what (if any) utility function a forager is maximizing.

## Materials and Methods

### Forager Dynamics

We consider a foraging animal subject to predation and moving through a continuous, two-dimensional domain Ω. We assume that the forager plans up until a finite, positive time *T*, which we call the planning horizon. Reasonable planning horizons will depend on the problem being studied, but we will generally choose *T* to be on the order of one day or one night. One reason for this is that we explicitly model spatial interactions between predators and prey through chases, with each chase lasting on the order of minutes. Thus, timescales on the order of seasons, while theoretically possible, will not generally be computationally tractable for our model. We track the forager’s location in Ω and its current energy level as functions of time, denoted **y**(*t*) and *ξ*(*t*) respectively.

As the forager moves through its environment, it may be spotted by a predator. We model this as a random event occurring at a rate *µ*_*s*_, i.e., the probability of the forager being spotted in a small time interval [*t, t* + *τ*] is approximately *µ*_*s*_*τ*. If a forager is spotted, we assume it is not killed instantly; instead, a chase ensues in which the forager may be killed or may escape. Each of these alternatives also occur at random times, governed by rates *µ*_*k*_ and *µ*_*g*_ respectively. We encode the current status of the forager relative to predators by assigning it one of two “operating modes:” foraging (mode 1) and escaping (mode 2). The current mode can thus be thought of as the state of a continuous-time Markov chain, illustrated in Figure 1a. (For convenience, we do not explicitly implement the absorbing mode 3, but simply set *ξ*(*t*) = 0 permanently whenever an animal dies from starvation or is killed by a predator.) If *η*(*t*) is a random variable that encodes the current state of that Markov chain, then the forager’s complete state at the time *t* is (**y**(*t*), *ξ*(*t*), *η*(*t*)). More generally, there is no reason why the rates must be constant, so *µ*_*s*_, *µ*_*g*_, and *µ*_*k*_ can be functions of the forager’s full state and time, though we only consider location-dependence in our examples.

**Figure 1:**
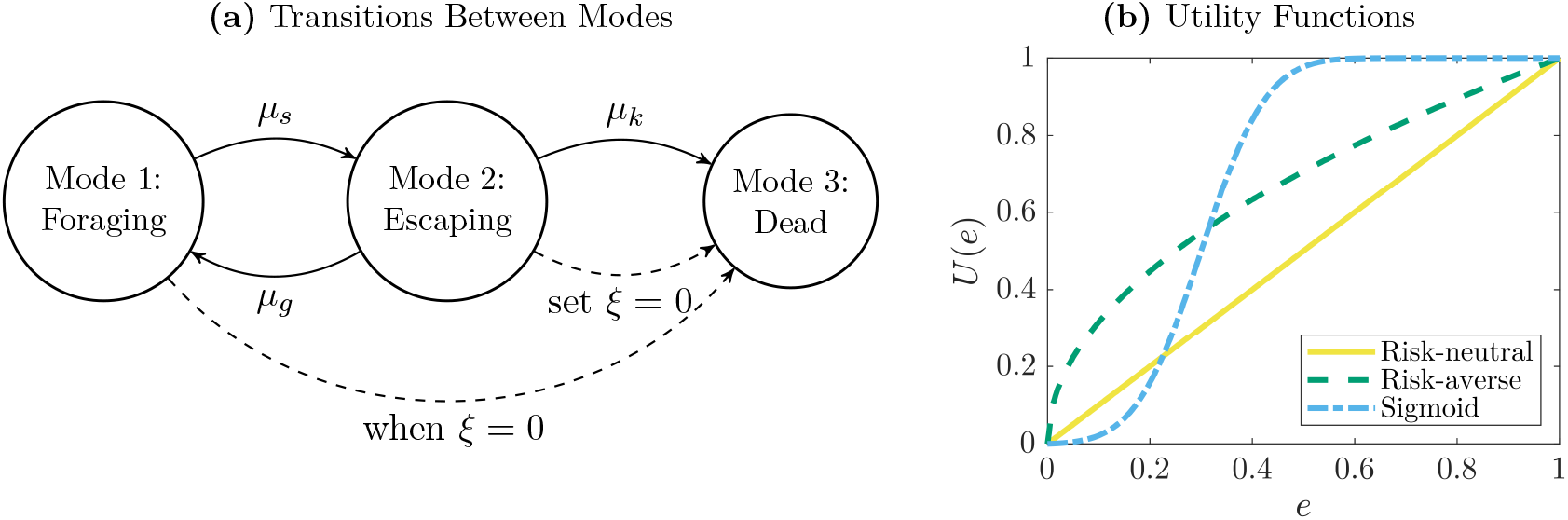
A forager maximizes its *expected* terminal utility due to random interactions with predators. **(a)** The forager’s current mode can be foraging (mode 1, actively collecting food), escaping (mode 2, moving in the immediate presence of a predator), or dead (mode 3). Each mode specifies its own dynamics for energy and position. Transitions between modes are stochastic and governed by known rates: *µ*_*s*_, the rate at which the forager is spotted by a predator; *µ*_*g*_, the rate at which a pursuing predator gives up on a chase; and *µ*_*k*_, the rate at which a pursuing predator successfully kills the forager. In general these rates may depend on time, the forager’s current energy level, and its current location, though our examples in the Results section consider location-dependence only. The forager deterministically transitions to mode 3 if it ever starves (reaches *ξ* = 0). **(b)** Possible functional forms for a utility function representing future fitness. Although utility is only a function of the forager’s final energy, the trajectory-dependence of the final state means that this function affects the optimal behavior at all times. Additionally, each utility function can be thought of as encoding specific risk preferences, discussed in detail in Section Utility Maximization.

Since the operating mode impacts how the forager interacts with its environment, we must equip each mode with its own location and energy dynamics. While the transitions between these modes are stochastic, we assume that the dynamics within each mode are deterministic, meaning that this model falls within the framework of piecewise-deterministic Markov processes (PDMPs) [Davis, 1984]. We use a simplified model of motion wherein the forager can change its direction of motion instantaneously, and the forager’s speed is determined by its state, but does not depend on the chosen direction^1^. Starting from some initial position **y**(0) = **x**, the forager’s position thus evolves according to

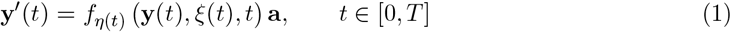

where *f*_*m*_(**x**, *e, t*) is a mode-dependent scalar speed function and **a** is a unit vector describing the chosen direction of motion. When we view this as an optimization problem, the animal will select a travel direction **a** as a function of its current state (position, energy, and mode) and time.

We assume that the forager continuously expends energy at a mode-dependent rate *K*_*m*_(**x**, *e, t*), and if the energy ever reaches zero the forager dies of starvation. In mode 2 (escaping), the animal cannot collect food, and thus when *ξ*(*t*) *>* 0 the energy dynamics are simply

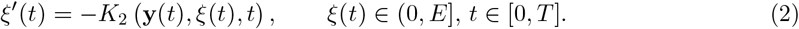

In mode 1 (foraging), the animal gains energy by collecting food from the environment at some rate *F* (**x**, *t*), which we compute as a weighted average of an underlying density of food *ψ*(**x**, *t*) in calories per unit area (see the Supplementary Materials A for details). In this case, the energy dynamics are

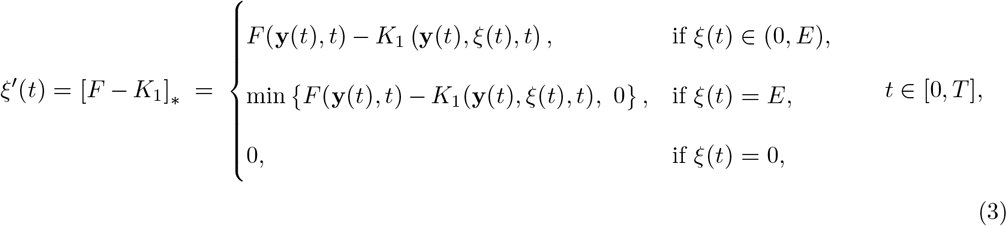

where the second case ensures that the forager never exceeds its maximum attainable energy level *E*, and the third case ensures that a forager who died from starvation or predator attack will not gain any energy in the future.

### Utility Maximization

We assume that the forager seeks to maximize a terminal reward, in our case the *expected* utility of its final energy level. The expectation accounts for the fact that interactions with the predator, and thus the forager’s final state, are stochastic. The most common interpretation of the terminal reward is as a measure of or proxy for future reproductive success (e.g., *fitness*) as a function of an animal’s state [McNamara and Houston, 1986, Mangel and Clark, 1986]. While not a requirement, many fitness functions depend on the animal’s energy level. Some functional forms that have been used in models of optimal foraging are step-functions [Rands, 2017], linear functions [Houston et al., 1993], [Fiksen et al., 1995], various concave functional forms [Houston et al., 1993, Sirot, 2019], sigmoid curves [Houston et al., 1993], [Caraco and Chasin, 1984], and occasionally convex functions [Kirby et al., 2000]. Additionally, [Brown and Kotler, 2004] propose various measures of fitness based on a combination of survival probability and energy gained from foraging.

We do not prescribe a single utility function, but instead consider three possible functional forms for the utility, and study their impact on optimal foraging behavior.^2^ We generally assume that an animal earns utility 0 if it is eaten. Throughout the paper, we compare three commonly used functional forms for utility shown in Figure 1b and given by

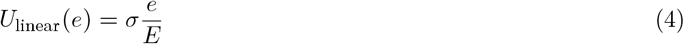

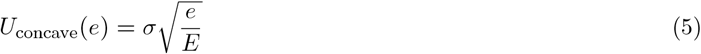

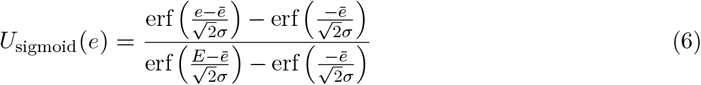

where *σ* is a shape parameter, *ē* is an energy threshold, and 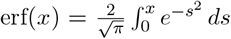 is the standard error function. The utility functions we consider can further be thought of as encoding risk preferences for a forager whose future fitness is proportional to its energy level after a foraging bout; i.e., here *e* = *ξ*(*T*) is a random variable, which is equal to zero if the animal dies earlier. In this framework, a risk-neutral planner has a linear utility (i.e., has no preference between two reward distributions having the same mean reward) and a risk-averse planner has concave utility (i.e., prefers distributions with lower variance, all else being equal). The sigmoid utility represents a planner that is risk-seeking at low expected utility and risk-averse at high expected utility. For steep curves (low values of *σ*) this approximates a step-function, the utility function associated with maximizing the probability of ending above a specific energy level. Though we focus on these three functional forms, our general approach is equally usable with a wide range of utility functions that depend on the forager’s state and time.

We use a dynamic programming approach to compute the policy that maximizes the expected utility. To do this, we assign each mode a value function *u*_*m*_(**x**, *e, t*) that represents the optimal expected reward-to-go for a forager in state (**x**, *e, m*) at time *t*. Formally, we define this as

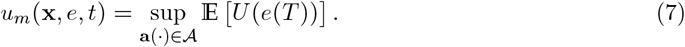

where the supremum (sup) is the maximum attainable (or almost attainable) expected utility. It can be shown that these value functions must solve a coupled system of Hamilton-Jacobi-Bellman PDEs (see the Supplementary Materials, B for a detailed formal derivation). Each term in each PDE describes a part of how the optimal expected utility changes with respect to time and the forager’s state. In both cases, all changes must sum to zero.

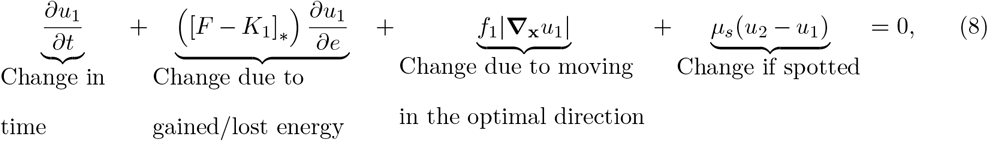

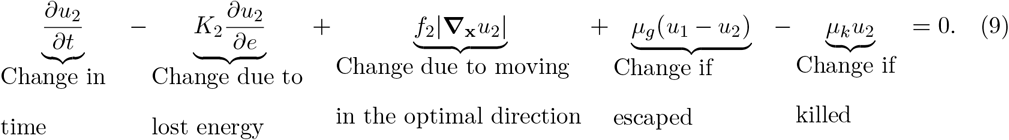

We use [*F* − *K*_1_]_*\**_ to ensure that the energy level stays between 0 and *E*, as defined in formula (3). We impose the boundary condition *u*_*m*_(**x**, 0, *t*) = 0, which ensures that the reward is 0 if the forager dies, and the terminal condition *u*_*m*_(**x**, *e, T*) = *U* (*e*), so that the forager collects a reward of *U* (*e*) if it survives the entire planning horizon. We solve this system backwards in time using an explicit finite difference discretization (outlined in detail in the Supplementary Materials, Section D).

## Results

### Balancing Risk and Reward

For our first example, we consider a forager that faces a trade-off between risk and reward. The environment is assumed to contain two food-rich regions (similar in structure to food patches in a discrete model), with the upper patch containing more food than the lower (Figure 2a). We further assume that the forager begins and ends its foraging bout within a refuge (outlined in white), which represents a burrow, den, or other area of increased cover. When in the refuge, the forager cannot be spotted or killed by a predator (*µ*_*s*_(**x**, *e, t*) = *µ*_*k*_(**x**, *e, t*) = 0) and has an increased chance of escaping (*µ*_*g*_ is increased by a factor of 10), but also cannot collect any food (*F* (**x**, *t*) = 0). Outside of the refuge, we choose the spotting rate *µ*_*s*_(**x**, *e, t*) so that the direct path to each food patch passes through an area of elevated spotting risk (Figure 2b). We will generally assume that this risk is higher along the path to the better food patch, with the animal aware of this *risk premium ρ* when it chooses which patch to visit.^3^ We use a constant kill rate *µ*_*k*_(**x**, *e, t*) = 4.0, give-up rate *µ*_*g*_(**x**, *e, t*) = 1.0, speed *f*_*m*_(**x**, *e, t*) = 2.0, and metabolic cost *K*_*m*_(**x**, *e, t*) = 3.0 in both modes (*m* = 1, 2). We allow the animal to reach a maximum energy level of *E* = 10 and assume the foraging bout lasts until time *T* = 1. Combined with our choice of *K*_*m*_, this implies that any animal that begins at an energy level *e*_0_ *>* 3 will not starve, even if it does not collect any food.

**Figure 2:**
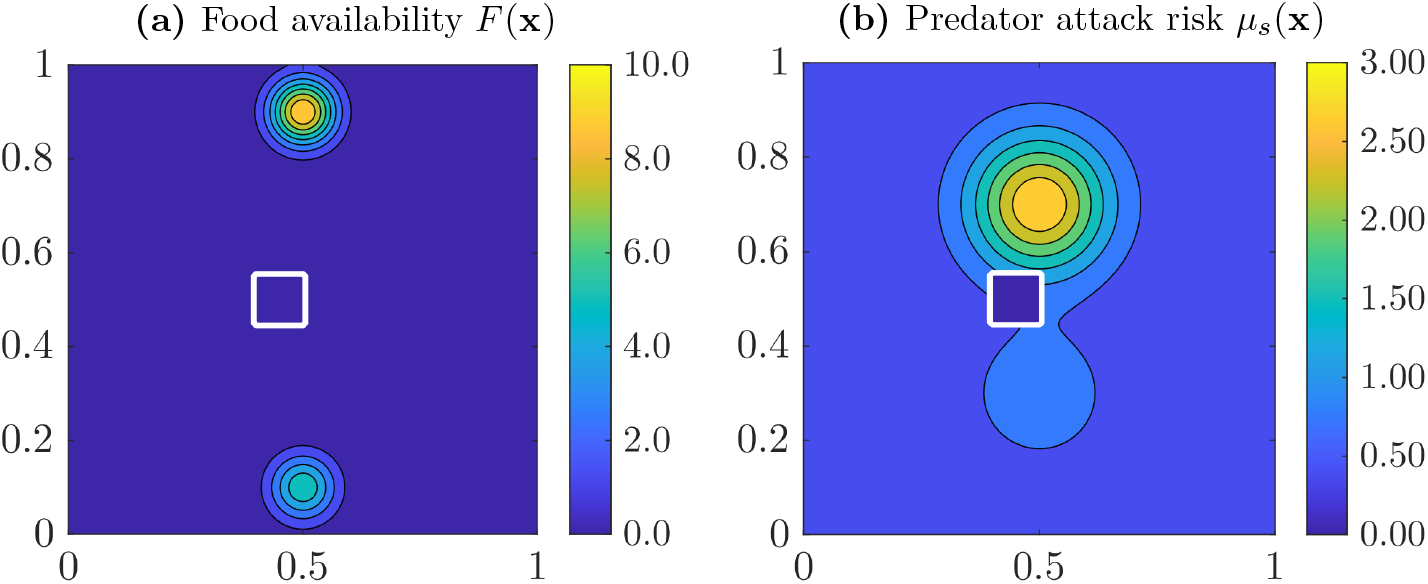
An inhomogeneous continuous environment similar in spirit to a discrete two-patch model. Food is concentrated in two small regions of the domain, where the upper is more lucrative (see the food availability *F* (**x**) in subfigure **(a)**) but also more risky (see the predator attack risk *µ*_*s*_(**x**) in subfigure **(b)**). In both panels, the white square represents the refuge, where the foraging bout must begin and end and where the forager can hide from predators without any risk but cannot collect food. Exact expressions for each quantity are provided in the Supplementary Materials E. The areas of highest risk lie along the most direct routes from the refuge to each feeding ground. This impacts the paths selected by a rational forager even when no predators are currently present. Both the choice of the feeding ground and the extent of a risk-avoidance detour that the forager is willing to take depend non-trivially on the initial energy, the planning horizon, the level of “risk premium” *ρ* (differentiating the upper and lower halves of the domain and defined formally in the text), and the forager’s utility function. These dependencies are further illustrated in Figures 3 and 4.

We focus on three questions: first, if the animal chooses to forage instead of staying in the safety of its refuge, which patch should it head to? Second, what is the optimal path to and from the chosen patch? Third, how does the optimal behavior depend on the shape of the forager’s utility function? While the first of these questions could potentially be answered using a “traditional” (discrete) foraging model, the other two questions leverage the continuity of our model to extract additional information about the optimal behavior.

Figure 3 summarizes how the preferred foraging patch changes as we vary initial energy level and risk premium for four possible utility functions—risk-neutral, risk-averse, and two versions of a sigmoid utility.^4^ For both sigmoid utility functions we set the threshold energy to *ē* = 0.3*E*, which means the forager can easily surpass the inflection point of the curve for most initial energy levels. We vary the steepness of the sigmoid curve, with the default curve having a shape parameter of *σ* = 0.1*E* and the wide curve having *σ* = 0.2*E*. The risk premium is varied by scaling the risk level in the upper half of the domain, while keeping the risk level in the bottom half the same across all simulations.

**Figure 3:**
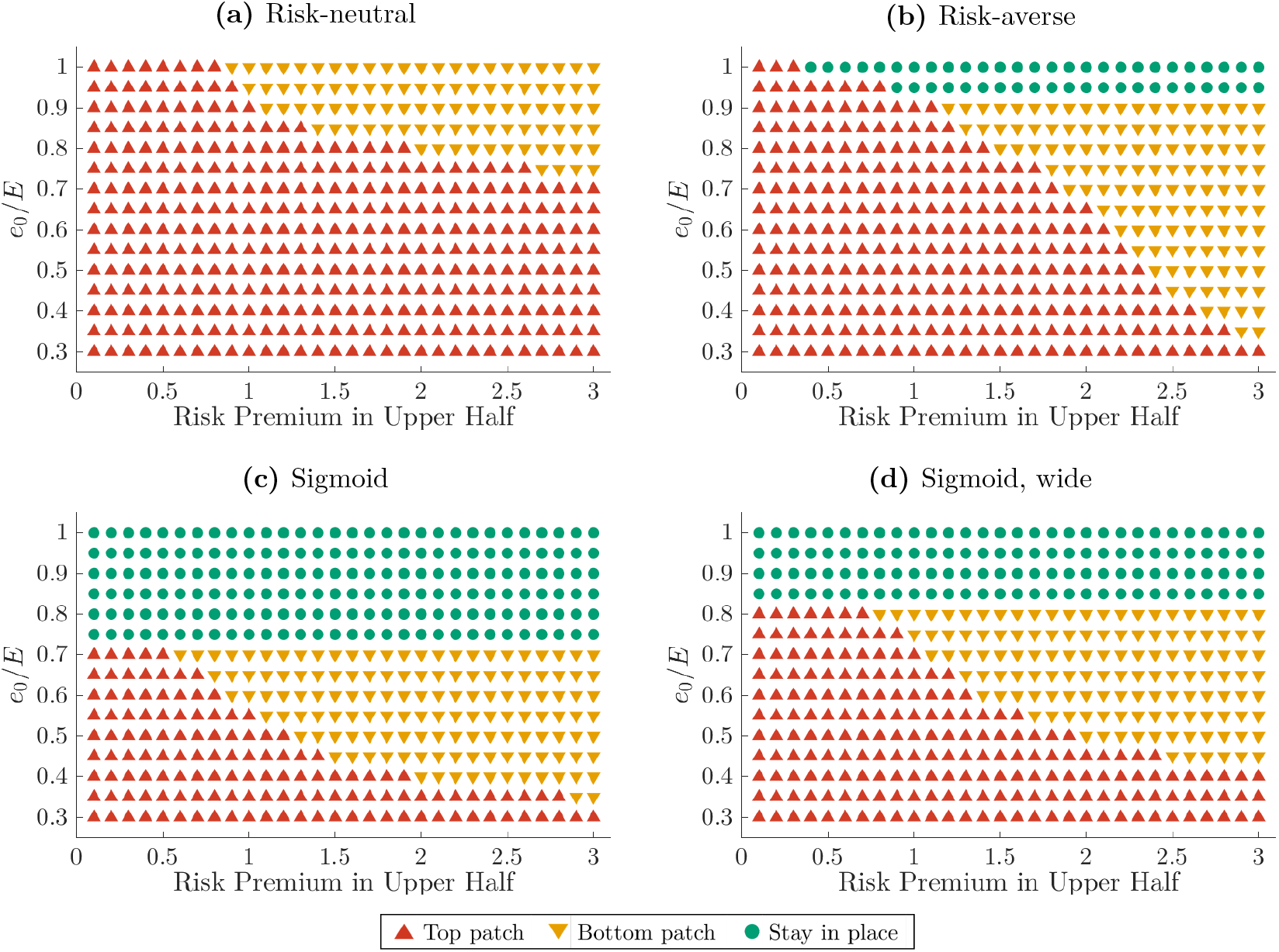
Preferred feeding ground for variable initial energy, risk premium, and utility function. Patch choice represents which food rich region the forager initially plans to visit (this plan may change due to the appearance of a predator). Across all utility functions, the forager is more likely to choose the riskier option when its initial energy is low (it is close to starvation), though this is mitigated by the level of the risk premium. There are significant regions of parameter space for which more than one utility function leads to the same patch choice, e.g., at low values of the initial energy and risk premium or for very high values of the risk premium. In these cases, it is natural to ask whether other aspects of the forager’s behavior differ across utility functions as this has implications for experiments that seek to infer risk preference from behavior. Figure 4 examines the forager’s planned *trajectories* for a range of initial energy levels.

In general, the risk-neutral planner is the least conservative in its choice of food patch, and the narrow sigmoid planner is the most conservative. The wide sigmoid planner makes qualitatively similar patch choices to the narrow sigmoid planner, but it has a higher baseline risk tolerance. Indeed, when the risk premium is high and the initial energy is low, the wide sigmoid planner is less conservative than the risk-averse planner. This is because the wider sigmoid curve more closely approximates a linear, or risk-neutral, planner for energy levels near the inflection point of the curve. However, because the curve is not truly linear, the wide sigmoid planner is still more conservative in its patch choice than the truly risk-neutral planner.

All planners except the risk-neutral one sometimes make the choice to simply stay in place rather than collecting food; in those cases, the guaranteed survival outweighs any possible reward associated with venturing out. Since we do not vary the risk level in the bottom half of the domain, any animal that prefers staying at home to visiting the lower patch will do so for all levels of the risk-premium. However, it is still possible that the top (more risky) patch will be preferred over both of those for some values of *ρ*, as can be seen in Figure 3b for the risk-averse planner. These preferences could have consequences for how often foragers with differing risk preferences leave safety to collect food. Animals that are more risk-averse may wait until they are at lower energy reserves to begin foraging relative to their less risk-averse counterparts.

While the shape of the utility function *U* clearly impacts the optimal patch choice, there are significant regions of parameter space where the same patch is optimal for several different utility functions. This is especially true at low levels of the risk premium and at the lowest levels of initial energy. In experiments that test risk-preferences for foraging animals, it is thus important that the risk be substantial enough to motivate different behaviors. It also suggests that examining a range of starting energy states is important for discerning the animal’s risk-preferences (as captured by the utility function).

However, the patch choice is not the full story of forager’s optimal behavior, as two animals exploiting the same feeding ground could follow substantially different routes to and from it. A longer detour bypassing high predator density areas can often decrease the overall risk, but will also decrease the time available for feeding. In balancing the time spent on feeding versus traveling, the forager must take into account the food patch quality, predator density variations, its level of hunger, and the utility function. Figure 4 shows how the shape of optimal trajectories changes with utility function and initial energy level *e*_0_ for a fixed value of the risk premium. In general, all foragers take safer (and longer) trajectories as *e*_0_ increases. Additionally, most utility functions predict trajectories that are similar on the outbound and inbound trips, with the exception of the risk-neutral planner, which chooses much shorter/direct routes on the way back. Such a strategy leads to higher expected utility, but also increases the variance in outcomes. For the same initial energy level and patch choice, the risk-averse planner takes longer detours than the risk-neutral planner on both legs of the trip. On the other hand, while the sigmoid planners do not always take the most conservative paths, they are the earliest to switch to the low-risk patch when the risk premium increases. As with the patch selection, the shape of chosen trajectories depends on the initial energy level of the forager, reinforcing the value of varying initial energy levels when discerning risk-preferences in field experiments. Overall, the observed routes to and from the feeding locations can provide valuable additional information for selecting a suitable utility function in models of foragers’ behavior.

**Figure 4:**
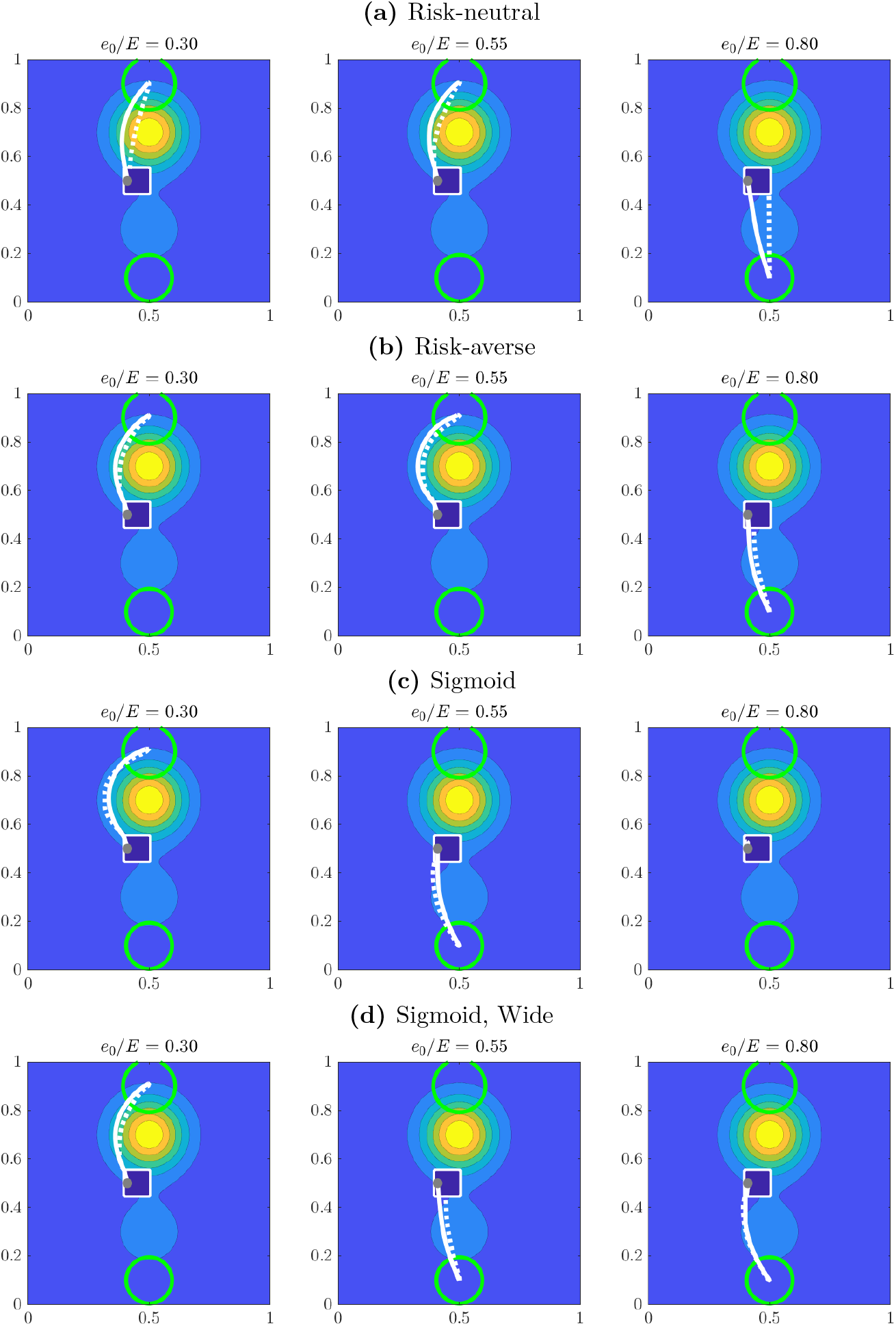
Preferred paths for selected initial energy levels and a fixed risk premium (*ρ* = 2). Paths to feeding grounds shown in solid white; return paths shown in dashed white. All subfigures present optimal trajectories chosen with possible predator encounters in mind—these are the paths followed if predators are feared but not really encountered along the way. As the initial energy increases, the forager chooses lower-risk options—longer detours, safer destinations, or even staying within the refuge. This matches the trend seen in Figure 3. Comparing different utility functions, foragers who choose the same feeding ground will sometimes take different paths to and from it, e.g., column 2 of **(a)** and **(b)** or column 1 of **(c)** and **(d)**). In those cases, risk preferences are only discernible if we consider factors other than patch choice.

### Foraging Samango Monkeys

We now consider an example based loosely on data from foraging Samango monkeys studied in [Coleman and Hill, 2014] and [Coleman et al., 2021]. Samango monkeys are small, arboreal primates that primarily forage for fruit in trees, travel in groups, and face predation from large birds of prey, snakes, and jaguars [Coleman et al., 2021]. One method that Samango monkeys use to avoid predators is to have a lookout, who will sound an alarm call if a predator is spotted, allowing the group to escape [Coleman and Hill, 2014]. While we do not model intra- or intergroup dynamics, we do assume that monkeys are aware of a predator when it appears and try to escape. In the study range, the primary predator encountered by the monkeys were eagles [Coleman and Hill, 2014], so we focus on just this predator-prey system. We track the position of the foraging monkeys in 2-dimensional space (though in reality they can also move up and down in the tree canopy).

We report our estimated model parameters in Table 2. All parameters are assumed constant except for food density *ψ*,^5^ spotting rate *µ*_*s*_, and give-up rate *µ*_*g*_. The relative magnitudes of *ψ* and *µ*_*s*_ were estimated using color data from images in [Coleman and Hill, 2014] and then linearly rescaled using the parameters in Table 2. We set *µ*_*g*_ to be complementary to *µ*_*s*_, i.e., *µ*_*g*_ is a linear rescaling of 1 − *µ*_*s*_/ max(*µ*_*s*_). Our versions of the resulting distributions are shown in Figure 5. The foraging area reported in [Coleman and Hill, 2014] is 0.547 km^2^ but for simplicity we assume that the map has a width of 1 km and a height of 0.7 km, which results in a comparably sized domain once obstacles are accounted for. When simulating forager trajectories, starting locations were chosen from areas identified as being close to sleeping sites in [Coleman and Hill, 2014], and the planning horizon was on the scale of a single day. To reflect the fact that most monkeys are likely far from starvation, we use an initial energy of *e* = 0.5*E* = 700 kcal which means that the monkeys cannot die of starvation within the time period we consider.

**Table 1:** Model parameters. We formulate a continuous model of position and energy and assume the forager selects its direction of motion in order to maximize its expected terminal utility (see Figure 1b). Food availability is computed based on a weighted average of the food density near the forager’s current location, and can be thought of as the rate of energy intake. The subscript *m* represents the forager’s possible operating modes (Figure 1a) and can take the values 1 or 2. We allow the metabolic costs and speed of travel to be mode-dependent; the animal’s full state (including mode) also influences its selection of the optimal direction of motion. As a result, the evolution of the forager’s energy and position is strongly impacted by the random mode switches.

| Parameter | Description |
| --- | --- |
| $\mu_s(\mathbf{x}, e, t)$ | Rate at which the forager is spotted by predators |
| $\mu_k(\mathbf{x}, e, t)$ | Rate at which a chasing predator kills the forager |
| $\mu_g(\mathbf{x}, e, t)$ | Rate at which a predator gives up on a chase (equivalently, the forager escapes) |
| $K_m(\mathbf{x}, e, t)$ | Metabolic cost |
| $f_m(\mathbf{x}, e, t)$ | Speed of travel |
| $\mathbf{a}_m(\mathbf{x}, e, t)$ | Direction of motion as a feedback control |
| $\psi(\mathbf{x}, t)$ | Underlying food density in the environment |
| $F(\mathbf{x}, t)$ | Food availability (rate of energy accumulation from foraging) |
| $U(e)$ | Utility representing future reproductive success |
| $\Omega$ | Domain |
| $E$ | Forager's maximum energy reserve |
| $T$ | Planning horizon |

**Table 2:** Parameter values and sources used for the stylized samango monkey example. Ranges are provided for parameters that vary across the environment, otherwise the parameter is assumed to be constant. The planning horizon of 6 hours roughly matches the amount of time samango monkeys spend traveling and foraging in one day [Coleman et al., 2021]. Food density, food radius, and harvest rate are used to calculate food availability *F* (**x**) (see Supplementary Materials Section A for details). When information directly pertaining to samango monkeys was not available, we used sources that study similar primates. While we aim to build a plausible model of the environment, we do not account for factors such as intergroup competition, the location of water, or other daily activities outside of foraging and traveling. We use the model to make qualitative predictions about the impact of the environment and utility function on foraging behavior (see Figure 6) rather than quantitative predictions about the trajectory of a specific animal.

| Parameter | Symbol | Range | Units | Justification/Source |
| --- | --- | --- | --- | --- |
| Metabolic cost | $K_m$ | 14 | $\frac{\text{kcal}}{\text{h}}$ | [Pontzer et al., 2014] |
| Speed | $f_m$ | 0.7 | $\frac{\text{km}}{\text{h}}$ | [Coleman et al., 2021] |
| Food density | $\psi$ | $2.00 \times 10^6 - 2.20 \times 10^8$ | $\frac{\text{kcal}}{\text{km}^2}$ | [Coleman et al., 2021] |
| Maximum energy | $E$ | 1400 | kcal | [Coleman et al., 2021],<br>[Zihlman and Bolter, 2015] |
| Planning horizon | T | 6 | h | [Coleman et al., 2021] |
| Spotting rate | $\mu_s$ | 0.01 - 0.12 | $\frac{1}{\text{h}}$ | [Coleman and Hill, 2014] |
| Kill rate | $\mu_k$ | 5.0 | $\frac{1}{\text{h}}$ | [Touchton et al., 2002] |
| Give up rate | $\mu_g$ | 1.0 - 11 | $\frac{1}{\text{h}}$ | [Cordeiro, 2003] |

**Figure 5:**
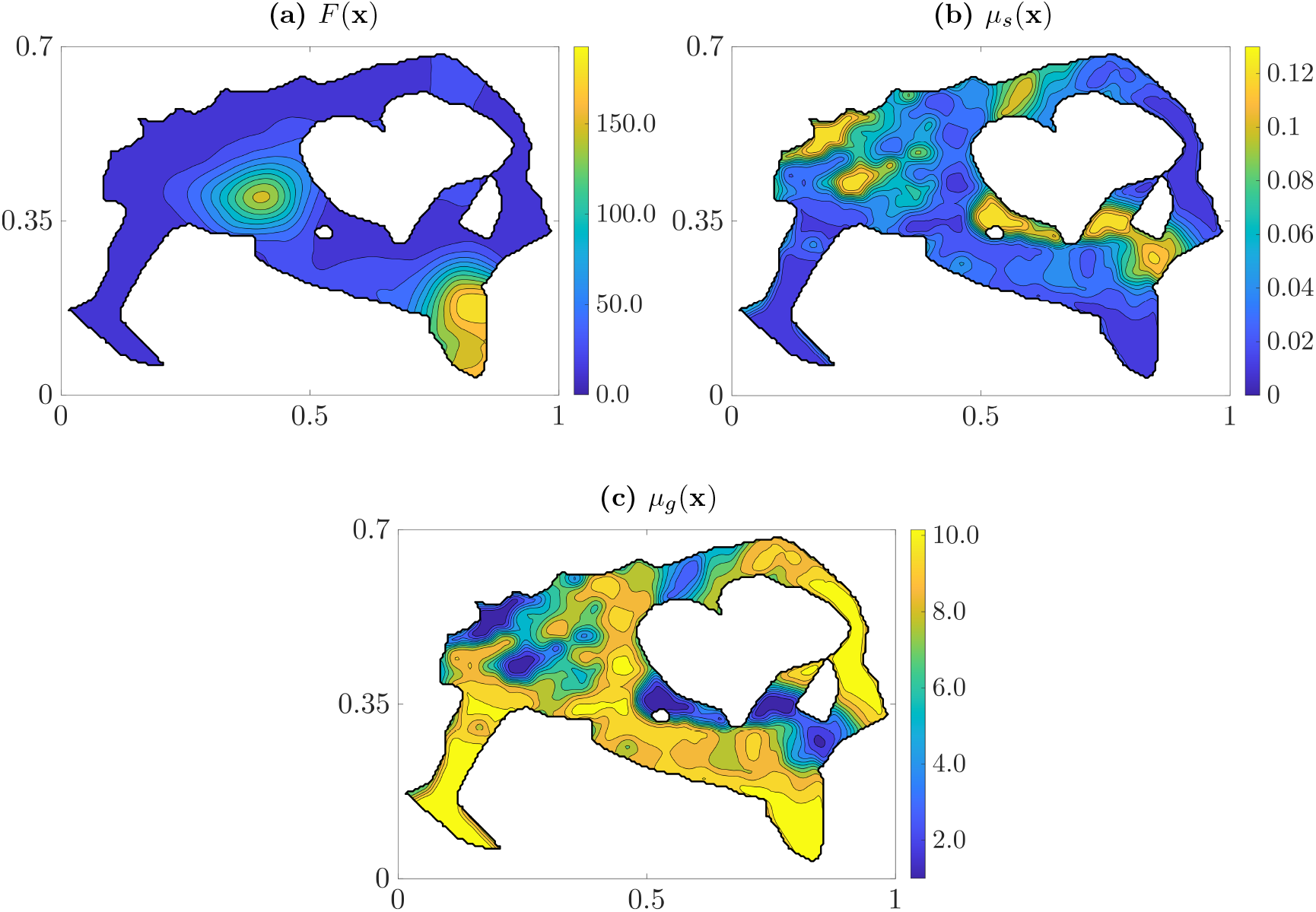
Stylized environment based on data for samango monkeys. Samango monkeys are small arboreal primates that mainly forage by collecting fruit in trees and are subject to predation from snakes, jaguars, and eagles [Coleman et al., 2021]. In [Coleman and Hill, 2014], the authors present detailed data about the monkeys’ environment in the form of heat maps. To create our stylized environment, we estimate the relative distribution of parameters using color data from the images and linearly rescale it using the parameters shown in Table 2. **(a)** Food availability is calculated based on food density (see Supplementary Materials Section A for details). Distribution of food density is based on Figure 1b in [Coleman and Hill, 2014]. **(b)** Distribution of predators is based on Figure 1g in [Coleman and Hill, 2014], which measures alarm calls due to eagles. We apply a smoothing filter due to small high-frequency variations in color data. **(c)** We choose to make the give-up rate complementary to *µ*_*s*_ (i.e., *µ*_*g*_ is a linear rescaling of 1 − *µ*_*s*_/ max(*µ*_*s*_)) because we assume that eagles are reluctant to pursue monkeys far from areas they normally hunt.

Figure 6 shows predicted optimal foraging trajectories for 1000 starting locations (chosen randomly from within identified sleeping sites) and the resulting habitat utilization. For all utility functions, optimal foraging paths avoid areas with high rates of predation even if those regions also have high food availability. This is in line with the main finding of [Coleman and Hill, 2014]: that predation risk was a consistent driver of habitat utilization. Additionally, our model predicts the formation of “channels” within which the optimal trajectories congregate. These channels are very similar for the risk-neutral and risk-averse utility functions, with most trajectories converging on a region in the lower-right corner of the domain that balances high food availability and low predation risk. In contrast, under a sigmoidal utility function, optimal trajectories favor areas of low predation risk essentially regardless of food availability. Here, we focus on the regime in which the monkeys’ energy reserves are safely above the threshold of starvation (and thus the inflection point of the utility function); as a result, they prioritize safety over taking risks for the sake of greater energy gain. For the initial energy level chosen, the habitat utilization map resulting from a sigmoid utility planner is clearly very different from those of risk-neutral or risk-averse planners. The habitat utilization by the latter two are much more similar, but some important differences are still discernible: e.g., the easternmost part of the domain has moderate utilization under risk-neutral planning and virtually no utilization under risk-averse planning. Predictions of this nature can help focus efforts to study risk-preferences on key areas of a habitat with largest expected differences in behavior.

**Figure 6:**
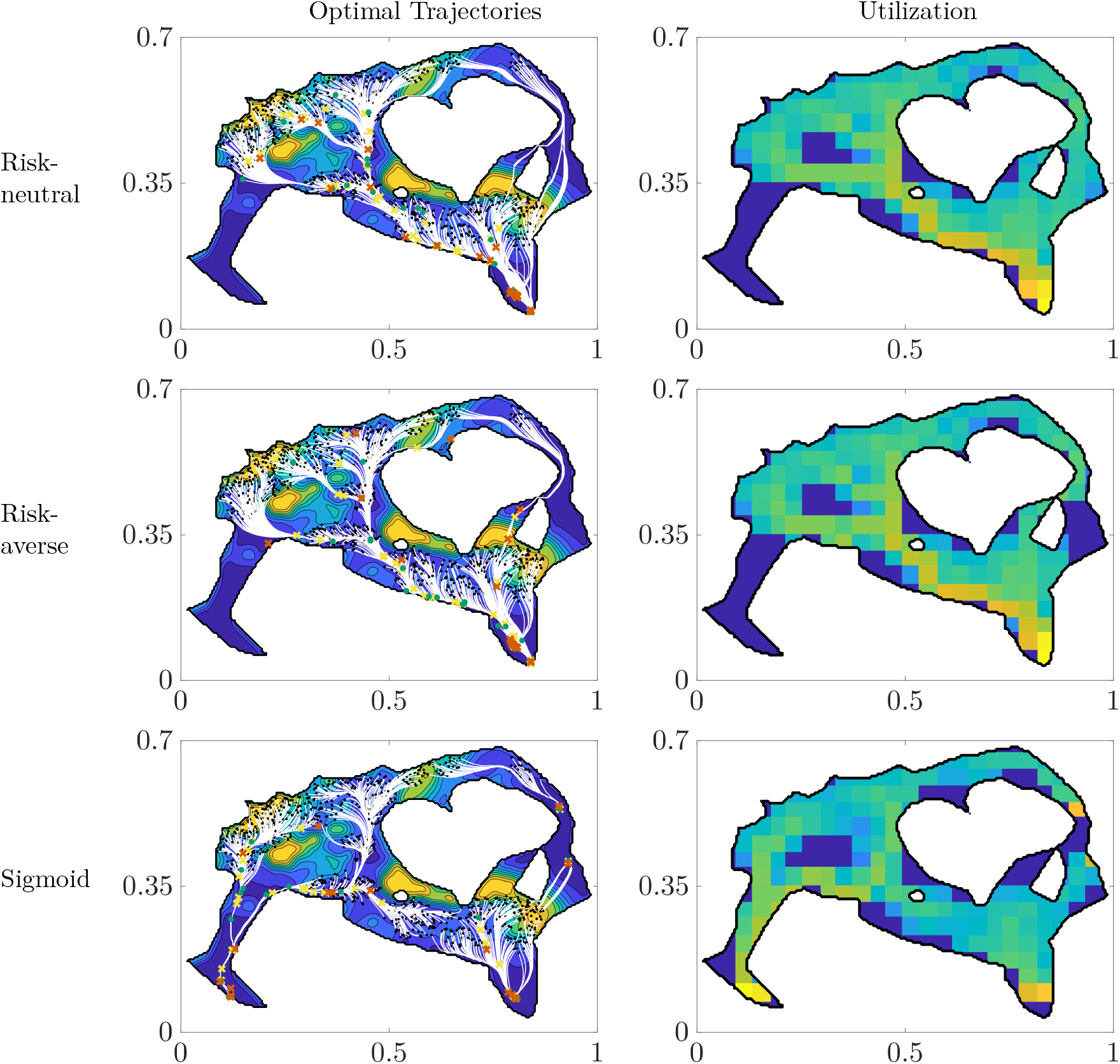
Monkeys’ optimal foraging trajectories and habitat utilization for three utility functions. Initial positions (black dots) are randomly chosen from areas identified as being close to sleeping sites in [Coleman and Hill, 2014]. Initial energy is set to *e* = 0.5*E* = 700, meaning that the monkeys cannot starve over the time horizon we consider. Left column: optimal trajectories and interactions with predators, where yellow “x”s are predator spotting events, green dots are successful evasions, and red “x”s are deaths. Trajectories congregate in “channels” as the monkeys head to their chosen destination. Risk-neutral and risk-averse planners generally end up at an area of high food availability, while sigmoid planners prioritize reaching nearby feeding grounds with low spotting rate. Right column: utilization is computed as the log of total time spent in a region divided by the area of that region. A notable similarity between our simulated utilization and the utilization estimated in [Coleman and Hill, 2014] is the dedicated avoidance areas with high risk of being spotted by an eagle. Our model is not sophisticated enough to reliably predict overall utilization but it provides qualitative insight into areas where we could expect large differences in monkey behavior based on the foraging objective.

Because we explicitly model interactions with predators, we can also examine predicted optimal behavior with and without predator encounters under each utility function. Figure 7 shows optimal trajectories followed from a single starting location under different utility functions if no predators are encountered along the way. These trajectories largely reflect the trends shown in Figure 6. Next, we consider two specific scenarios with predator interactions: (1) the monkey is immediately spotted by an eagle and (2) the monkey is spotted by an eagle after 0.5 hours. Even though the sigmoidal utility function leads to very different behavior in the absence of a predator, all planners take the same escape route if they are spotted immediately (Figures 8a, 8c, and 8e.) On the other hand, even though the risk-neutral and risk-averse planners have nearly identical trajectories in the absence of a predator, they take different escape routes when spotted at 0.5 hours (Figures 8b and 8d). Thus, the arrival of a predator can cause distinct planners to converge upon a highly favorable escape route, but can also cause otherwise very similar planners to take drastically different escape measures.

**Figure 7:**
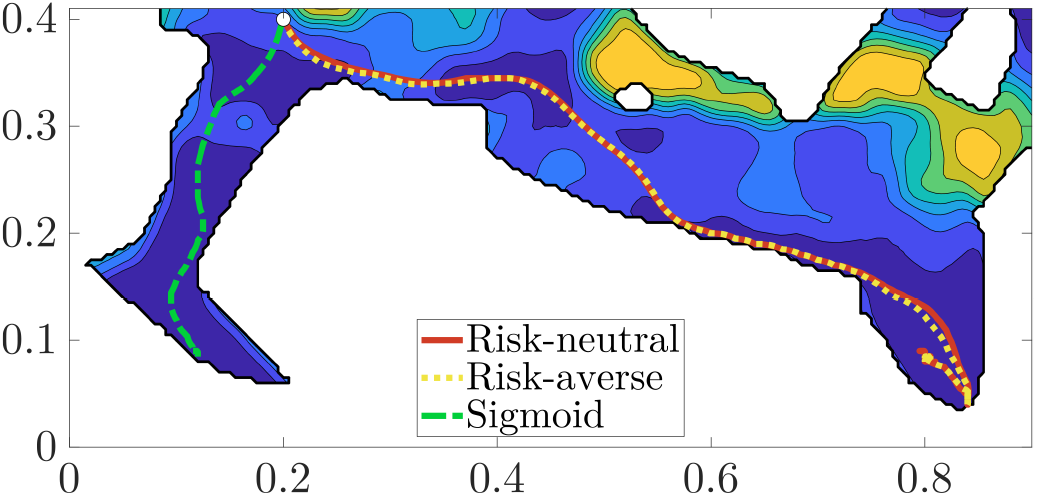
Sample “no predators encountered” trajectories under different utility functions. A monkey moves from the white dot to a chosen destination, where it spends the remaining time until the end of the planning horizon. Even though the monkey starts and remains in mode 1 (no predators), the possibility of encountering eagles still influences the chosen trajectory. The risk-neutral and risk-averse planners choose a feeding ground destination and follow very similar paths to it. (However, see Figure 8 for how differently they would handle an encounter with predators along the way.) Since the monkey begins above the energy threshold of the sigmoid curve and cannot fall below it in the time period we consider, the sigmoid-planner does not prioritize food as much and heads for the safest nearby location instead.

**Figure 8:**
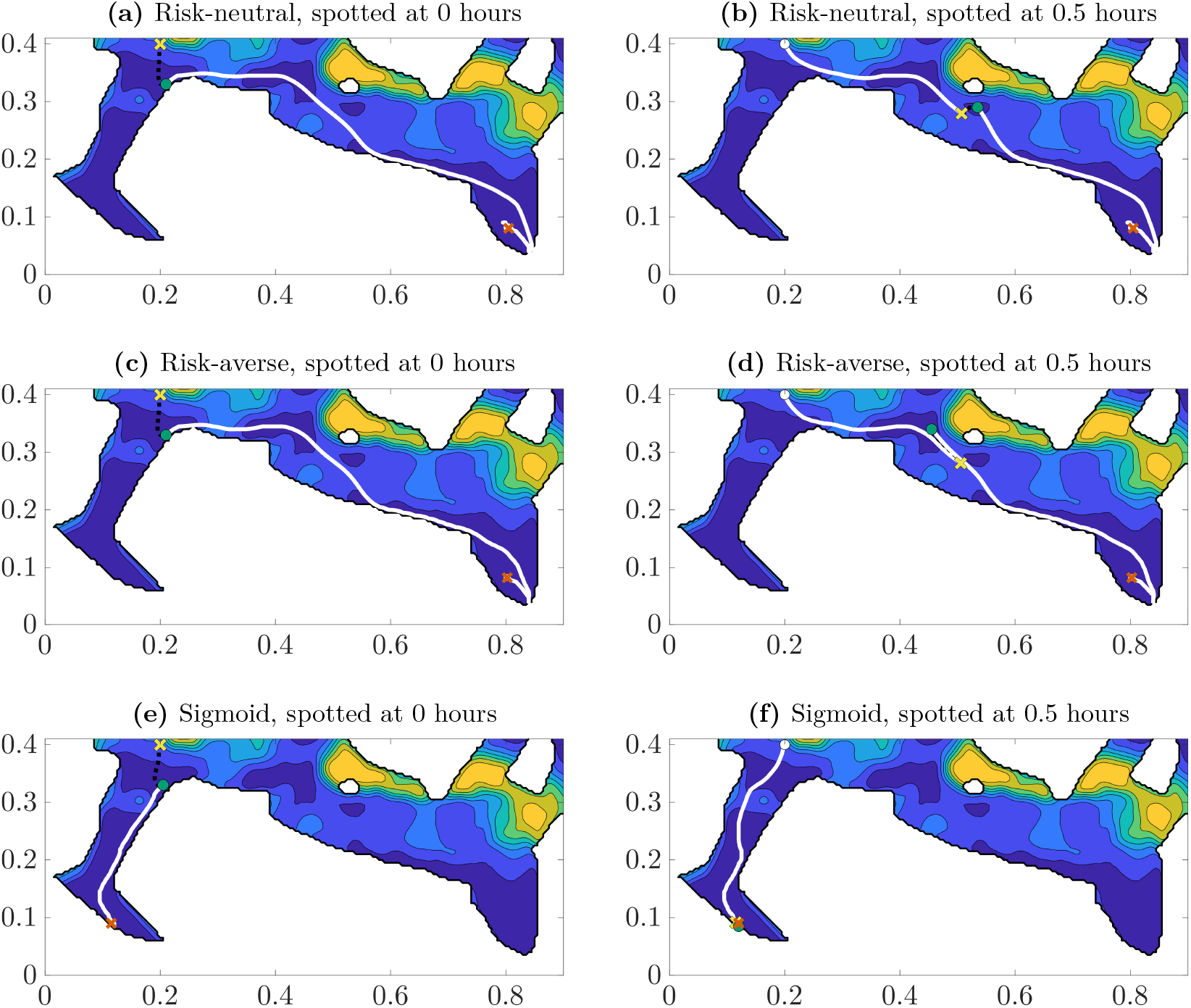
Sample trajectories with predator encounters across utility functions and spotting times. A monkey begins in the same starting location as Figure 7 but now it is spotted by an eagle either immediately (left column) or after 0.5 hours (right column). Yellow “x”s represent predator spottings and green dots represent successful escapes. In this scenario a monkey that is spotted immediately will follow the same escape trajectory (black dotted line), regardless of its utility function. Conversely, the risk-neutral and risk-averse planners (which follow near identical paths in the absence of a predator) take significantly different escape routes if spotted at 0.5 hours (subfigures **(b)** and **(d)**). The risk-neutral planner takes a slight detour to a safer location but does not greatly deviate from its planned path, while the risk-averse planner significantly backtracks along its previous route in order to escape. Here, we observe that monkeys that behave similarly in the absence of a predator may not react to a predator appearance the same way and vice versa.

## Discussion

Fear of predation has consequences at all scales in ecology. Exposing snowshoe hares to simulated predators during gestation reduced female survival by 30% and survival to weaning of offspring by more than 85% [MacLeod et al., 2018]. In general, the traits and actions evoked by predation risk have on average larger demographic impacts on prey populations than the actual deaths due to predation [Preisser et al., 2005]. Fear-induced responses by many individuals in a population can collectively affect community and large-scale ecosystem properties [Pinti et al., 2022]. For example, simulated presence of large predators reduced raccoon foraging in an intertidal zone community, in part by causing raccoons to spend more time foraging elsewhere. Reduced raccoon foraging resulted in 59% to 97% increases in the abundance of their main prey species; moreover, it also affected the survival and abundance of those raccoon prey species’ own prey and competitors [Suraci et al., 2016].

Mobile animals often try to reduce the predation risk (real or perceived) through their choices of feeding areas and movement paths while foraging (e.g., [Gilliam and Fraser, 1987, Lima and Dill, 1990, Brown and Kotler, 2004, Laundré et al., 2001, Brown et al., 1999, Coleman and Hill, 2014]). Optimal foraging models that incorporate predation risk have generally assumed discrete foraging “patches” varying in food availability and predation risk. The time resolution of such models is typically coarse and the physical proximity of patches is often ignored. An animal is essentially “teleporting” to and from its feeding grounds. while the predation risks en route to and at the chosen patch are lumped together. In contrast, the models introduced here treat foraging as a two-dimensional navigation problem in a complex, continuous landscape with food and risk varying continuously, and the forager’s behavior explicitly changing whenever a predator is encountered. Using stochastic dynamic programming, we find the optimal behavior for all state configurations, thereby predicting where an animal should choose to feed, what paths it should take to and from those locations, and how it should move when trying to escape a predator.

Understanding better how foragers decide where to go and what to eat is not just scientifically interesting – it is essential for predicting how these populations may be impacted by habitat modifications and changes in the abundance or types of prey or predator species. While we can often directly observe *what* a forager chooses to do in different circumstances in the real world, its decision algorithm has to be inferred indirectly from the observed behaviors. Our theoretical and empirically motivated case studies show that a forager’s chosen movement paths provide valuable information about its motivations, above and beyond what we could learn from just observing where it chooses to feed. In particular, these paths could be used to evaluate how the forager values different possible energy levels at the conclusion of a foraging bout (Figure 1b).

For any assumed utility function, the dynamic programming solution makes it easy to numerically construct a “phase diagram” of how foraging behaviors change as a function of the forager’s initial energy state and initial locations. Knowing which conditions lead to qualitatively different behaviors for different utility functions, one can identify which field experiments will be most informative about the unobserved decision-making process.

The model presented above assumes that a single forager does not have much impact on food availability in the environment. Situations where this impact is actually substantial are much harder to treat in our optimization framework – the food map changing due to forager’s behavior becomes a part of the (now, much larger) state space for making future choices. The resulting “curse of dimensionality” makes the direct use of dynamic programming computationally infeasible. To address this, we propose a “suboptimal foraging model” [Railsback, 2022], in which the forager plans over a short time horizon but periodically updates its plans based on how it has impacted the environment; see the detailed discussion in Supplementary Materials Section C. While this approach requires a careful selection of the frequency of such updates, we believe this is a useful step beyond modeling a single forager in an unchanging resource landscape.

One limitation of our model is the assumption of deterministic energy and location dynamics (Equations (1) through (3)). More generally, food uptake could be modeled as a series of random encounters with food items at a rate determined by the local food density, and death from starvation could be similarly treated as random (with an increasing probability as the energy level approaches zero). Randomness in movement paths could be modeled by a stochastic differential equation with additive Brownian motion reflecting an “exploration” aspect of foraging in addition to the “exploitation” already included in our model. Including these additional stochastic features would change the form of the governing Hamilton-Jacobi-Bellman PDEs, but the general numerical approach for solving it would be quite similar [Fleming and Soner, 2006].

Our current approach also does not allow us to consider effects of interactions between animals [Agrawal et al., 2007], with or without food depletion. One possible method within a continuous framework would be to model large numbers of interacting individuals using meanfield games (MFGs) [Huang et al., 2006, Lasry and Lions, 2007]. Multi-population MFGs can be also used to model complex cooperative interactions within cohesive groups and the competitive inter-group interactions of many animals exploiting depletable food resources [Coleman, 2013, Coleman and Hill, 2014]. Furthermore, using approximate dynamic programming methods designed to cope with high-dimensional state spaces will also expand the range of models that are computationally feasible [Powell, 2007].

A broader limitation of our modeling framework is the assumption that animals are perfect rational optimizers. In reality, there are reasons to believe that foragers would not have evolved perfectly optimal strategies since those rules would not necessarily provide large selective advantages over simpler decision rules [Janetos and Cole, 1981]. In fact, there is some evidence that simple, “good enough” rules produce behavior that is more closely aligned with reality [Crowley et al., 1990]. Thus, the trajectories presented here should not be thought of as quantitative predictions of animal behavior, but rather as qualitative descriptions of what a perfect optimizer would do. These are useful for two reasons: providing a bound on how well an animal could do in practice and providing qualitative insights for the behavior of “somewhat rational” and “somewhat informed” optimizers. One extension that could more accurately characterize trade offs between maximizing utility and relying on strategies that are simple and broadly applicable is multi-objective optimization, where a forager seeks to optimize two or more objectives at once. Efficient methods also exist for recovering such “Pareto optimal” solutions in the framework of dynamic programming [Mitchell and Sastry, 2003, Kumar and Vladimirsky, 2010, Désilles and Zidani, 2019].

Overall, we hope that this paper will motivate further work on modeling how mobile organisms choose to move in a landscape full of opportunities and threats. As animal tracking and biologging capabilities continue to improve, [Tomkiewicz et al., 2010, Kays et al., 2015], there are in-creasing opportunities to track animals over long periods of time and characterize qualitative relationships between animal state and behavior. These observations can in turn be compared to the optimal decision rules under different assumptions to learn more about how foragers trade off risks and opportunities. The consistently mediocre performance of species distribution models that seek to predict organism presence/absence or abundance using climate and habitat covariates (e.g., [Lee-Yaw et al., 2022, Liu et al., 2020]) suggests that mobile animals are not just “going where the climate suits my clothes” [Whitter, 1924]. Predicting how distribution and abundance will change in response to human impacts (either detrimental or intended to be helpful) will require a deeper understanding of how animals decide what to do where and when.

## Author Contributions

All authors contributed to the formulation of the mathematical model. SPE contextualized the findings and guided the modeling along with AV, who also guided numerical implementation. MG, SJ, NR, and AS developed the solution code. MG, SJ, NR, AS, and NGG contributed to the visualizations. MG, AS, and NGG contributed to data collection and processing. NGG, SJ, NR, and AS developed and performed initial numerical experiments, and MG, SJ, AS, AV, and SPE developed and refined the numerical examples included in the paper. MG, NGG, SJ, and AS were responsible for early drafting of the paper and results. MG, AV, and SPE revised and developed the manuscript, and all authors contributed to final editing and approval.

## Acknowledgments

This research was partially funded by the National Science Foundation (awards DMS-1645643 and DMS-2111522) as well as the Air Force Office of Scientific Research (award FA9550-22-1-0528).

## Data and Code Availability

All data and code used to generate the figures in this manuscript are available online at https://github.com/eikonal-equation/Navigating_LoF/tree/main.

## A Calculating Food Availability

Here we describe the details of how we compute food availability *F* (**x**, *t*), which can be thought of as the forager’s rate of energy accumulation. Within our domain Ω, there may be terrain in which the forager cannot collect food, such as obstacles (representing lakes or unnavigable terrain), or refuge areas (representing burrows or dens). The region in which the forager *can* collect food is called “forageable” and we denote it as Ω_*F*_. We assume a forager located at **x** collects food in a small circle of radius *r, D*_*r*_(**x**), and model the total food collected per unit time as proportional to a weighted average of the harvestable food throughout *D*_*r*_(**x**). When *D*_*r*_(**x**) is fully contained within Ω_*F*_, we have

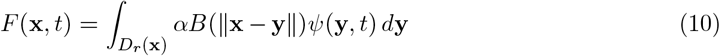

where the harvest rate *α* represents how efficiently the animal can access food resources, and *B* : ℝ *→* ℝ_+,0_ is a kernel function representing how the animal distributes its effort, with 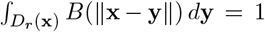 and *B*(*s*) = 0 for any *s > r*. More generally, if *D*_*r*_(**x**)⊄ Ω_*F*_, the animal spreads the same effort over a smaller area, *D*_*r*_(**x**) *∩* Ω_*F*_. To account for this, we introduce a normalizing factor

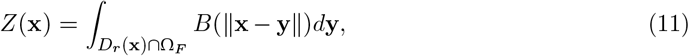

and derive a new expression for the rate of food accumulation:

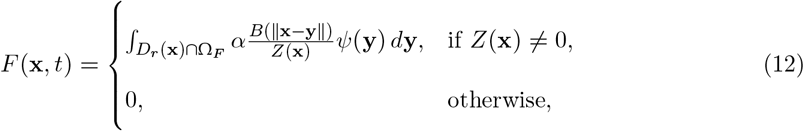

where we note that *Z*(**x**) = 0 only occurs if there is no forageable terrain in the area in which the forager collects food, and thus no food is obtained. All examples included in our paper use a constant function 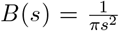 on *D*_*r*_, but our overall framework is more flexible. Another reasonable choice for *B* could be a truncated and normalized Gaussian, with the forager’s efforts more concentrated around its average location **y**(*t*).

## B Deriving the Governing HJB Equations

We use dynamic programming to derive the governing HJB equations, which determine the optimal policy for the foraging animal in each mode. For notational convenience, we will use **w** = (**x**, *e, t*) *∈* **W** = Ω *×* [0, *E*] *×* [0, *T*] to denote a generic position-energy-time triple, and ***ω***(*t*) = (**y**(*t*), *ξ*(*t*), *t*) to denote the current state of the animal with combined dynamics 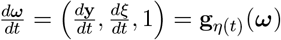. We now introduce the feedback control 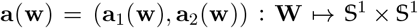, which prescribes the direction of motion in each mode as a function of the current state. (An animal that adopts policy **a**(·) must use **a**_1_(·) when it is in mode 1 and **a**_2_(·) when it is in mode 2.)

Let *J*_*m*_(**w, a**(·)) be the expected remaining reward for an animal in mode *m*, at state **w**, and using policy **a**(·). We can write an expression for *J*_*m*_ in each mode by explicitly computing the expected reward over the distribution of mode switches. Let *T*_*s*_ *> t* be a random variable denoting the first time the animal is spotted by a predator after some time *t*, and recall that the animal plans up until some fixed time *T*. Mode 1 ends: at *T*_*s*_, if the animal is spotted for the end of the time horizon (*T*_*s*_ *< T*), at any time when *ξ*(*t*) = 0 (representing death from starvation), or at *T* if neither of those events occurs first. We thus have the following expression for *J*_1_:

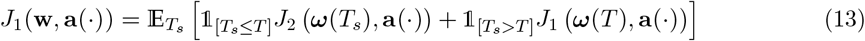

along with the boundary conditions *J*_1_(**x**, 0, *t*, **a**(·)) = 0 to represent death by starvation.

Similarly, define *T*_*g*_ and *T*_*k*_ to be the next times a forager chased by a predator manages to escape or is killed, respectively. Mode 2 ends when the predator gives up on the chase (*T*_*g*_ *≤* min{*T*_*k*_, *T*}), the forager is killed (*T*_*k*_ *≤* min{*T*_*g*_, *T*}) or dies from starvation (whenever *ξ*(*t*) = 0), or if neither of those events occurs and the forager survives until the end of the planning horizon *T* in mode 2 (min{*T*_*g*_, *T*_*k*_} *> T* and *ξ*(*t*) *>* 0 for all *t ≤ T*). Thus, we have the following expression for *J*_2_:

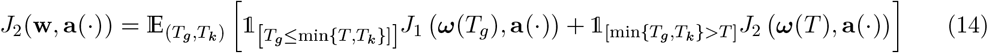

with a similar starvation boundary condition *J*_2_(**x**, 0, *t*, **a**(·)) = 0. We don’t specify *J*_3_ for mode 3 because when the animal is dead the remaining reward is identically equal to 0. That is also the reason why equations (13) and (14) do not contain terms explicitly associated with switching to mode 3: any such terms are equal to 0. We will also sometimes impose an additional constraint that the animal must be located in the refuge at time *T*, i.e., **y**(*T*) *∈* Ω_*R*_ where Ω_*R*_ is the set of points contained in the refuge. In those cases, we use the terminal condition 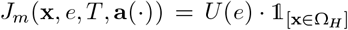 in both modes. For any areas through which the animal cannot travel, such as obstacles or points outside the domain, we impose the boundary condition *J*_*m*_(**x**, *e, t*, **a**(·)) = 0. Note that while we do not impose distinct terminal conditions for different modes, this would also be easy to handle with minimal changes.

Let *u*_*m*_(**w**) be value function for a forager in mode *m*, which represents the optimal expected reward-to-go for an animal at state **w** in mode *m* and can be obtained by maximizing *J*_*m*_:

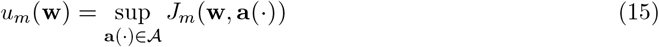

where 𝒜 is the set of measurable functions from *W* to 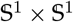. We now use Bellman’s Optimality Principle to derive a system of governing PDEs for *u*_1_ and *u*_2_. Consider a small time interval *τ <* (*T* − *t*) and assume that no mode switch occurs in [*t, t* + *τ*). The value functions must then satisfy the following optimality conditions:

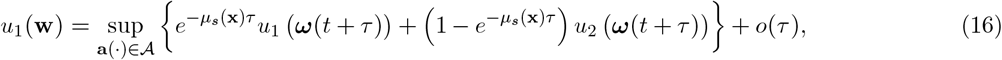

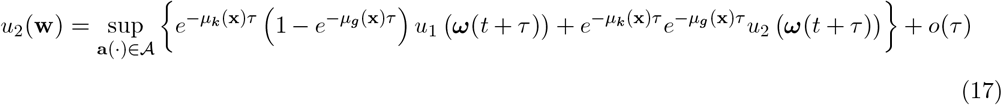

where for a random event governed by rate *µ* we have approximated the probability that the event occurs over *τ* units as 1 − *e*^−*µτ*^, and ***ω***(*t* + *τ*) represents the animal’s state *τ* time units after starting from ***ω***(*t*) = **w** and using control **a**(·). We next perform a Taylor expansion in *τ*. For the value function in mode 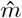 and for an animal that is in mode *m* at time *t* we have

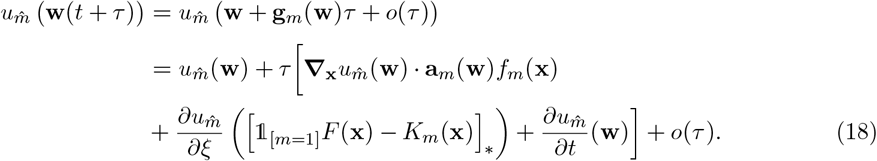

Substituting (18) into equations (16) and (17) and suppressing arguments, we find that the HJB PDEs for each mode are:

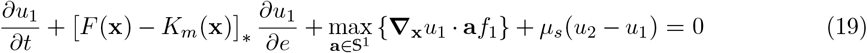

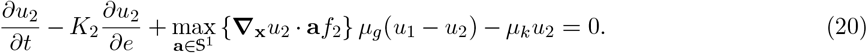

Using the fact that the maximum is attained with **a**_*m*_(**w**) = ***∇****u*_*m*_(**w**)/∥***∇****u*_*m*_(**w**)∥, we obtain equations (8) and (9):

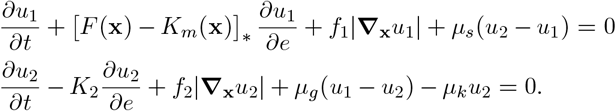

Finally, the boundary and terminal conditions for the value functions come directly from those for *J*_*m*_: *u*_*m*_(**x**, 0, *t*) = 0 and *u*_*m*_(**x**, *e, T*) = *U* (*e*).

## C Modeling Food Depletion

Throughout the main article, we assume that foragers do not deplete food as they exploit their environments, as in the classical optimal diet model. This is a reasonable assumption when resources are abundant or regenerate quickly. However, some foragers meaningfully deplete resources as they exploit them, and must account for this in their foraging decisions. Failure to account for this in a model can lead to unrealistic predicted behaviors. A forager that experiences food depletion but does not plan for it will generally achieve a lower energy level than it initially predicted, and may adjust its path in order to compensate (Figure 9). In principle, food depletion can be addressed by including the remaining food at every location as additional state variables in the dynamic programming model. Unfortunately, this approach quickly becomes computationally infeasible due to the “curse of dimensionality”—the computing time required to calculate the value function grows exponentially with the dimension of the state space. At the same time, it is disputed whether it is accurate to model foraging animals as making highly accurate long-term predictions about their environment [Railsback, 2022], so it is not obvious that we should model animals as exactly accounting for food depletion during an entire foraging bout.

**Figure 9:**
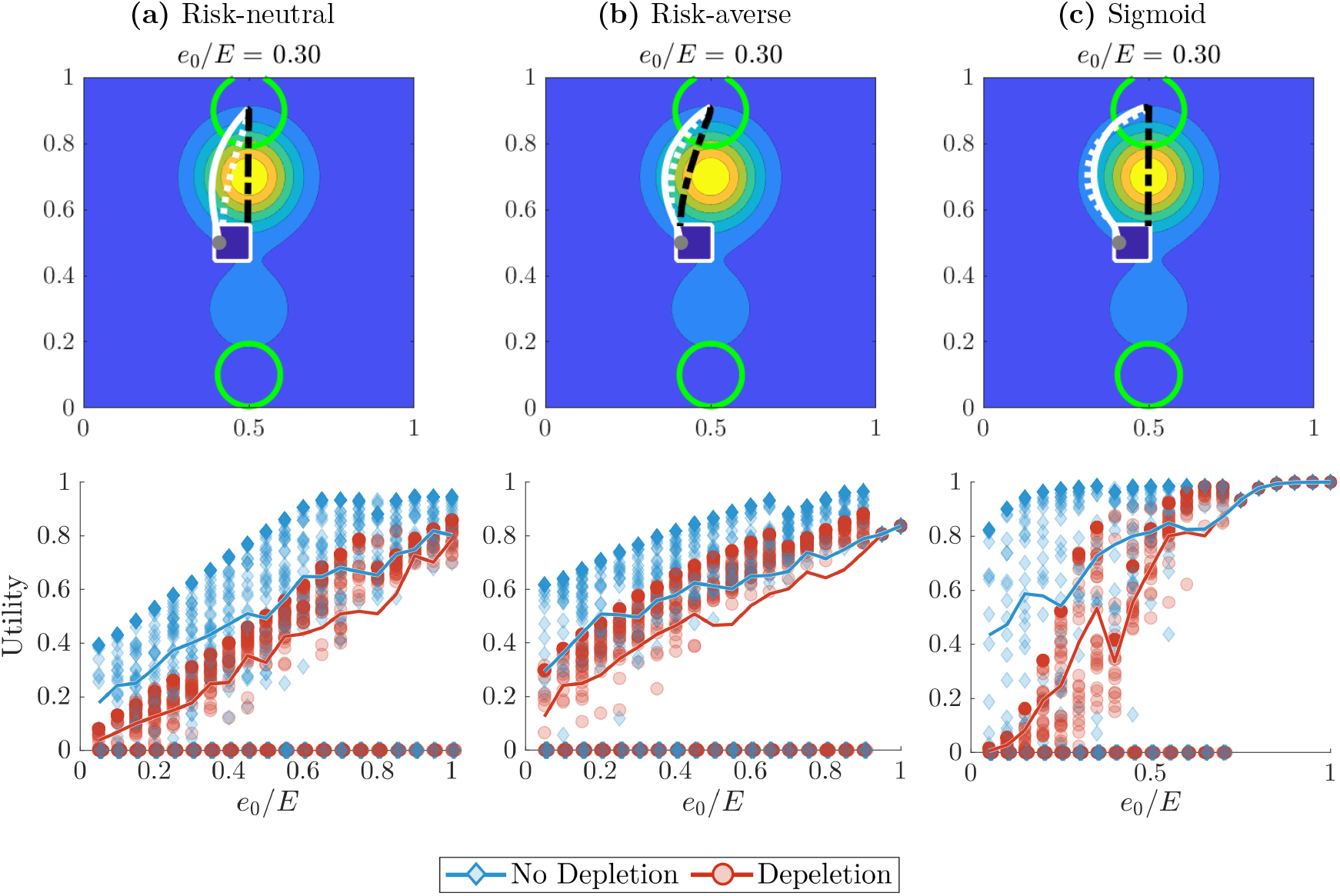
The impact of food depletion on a rational forager that *does not plan for it*. (Except for food depletion and initial energy levels, the setting is the same as in Figures 2 and 4 of the main manuscript.) When the forager depletes food, it receives less energy than it planned for and spends longer at the feeding ground to compensate. It subsequently takes shorter, riskier return trips to the refuge (top row, black dot-dash line) than the ones it originally planned (white dotted lines) without accounting for food depletion. Outbound trips (solid white line) are identical because no depletion has occurred yet. The bottom row shows realized terminal utilities with (orange circle) and without (blue diamond) food depletion, which vary for fixed *e*_0_ due to random interactions with predators. A forager that dies via starvation or is killed by a predator receives a utility of 0 (points at *U* = 0 have been horizontally perturbed to indicate frequency). As expected, utility is on average lower when the forager depletes food (orange and blue solid lines). Overall, food depletion may significantly impact the forager’s optimal trajectory and final utility in a way that is nonuniform across utility functions and initial energy levels. However, exactly accounting for the impact of food depletion on an optimal forager is computationally infeasible and may not be ecologically sound. We instead propose a model in which the forager makes inaccurate short-term predictions about its environment, but periodically updates them to account for the food it has depleted.

Motivated by these considerations, we now describe a preliminary “suboptimal” foraging model in which a forager makes decisions based on imperfect, short-term predictions for the future state of its depleting environment. [Railsback, 2022] outlines a “suboptimal foraging” approach for individual based models (IBMs) where foragers evaluate the quality of a patch under the assumption that its state remains constant over a long period of time. We instead assume that the animal makes myopic estimates and decisions over a *short* time period (stage) relative to the duration of a foraging bout. Specifically, we suppose that a forager makes plans to maximize its expected utility *S* time units later, on the assumption that food availability will not change from its current state. It then behaves accordingly for *S* time units, collecting and depleting food as it moves. At the end of each short stage, the forager becomes aware of the updated map of food availability, and makes new plans for the next short stage. This continues until the end of the time horizon *T*. This “multistage model” converts the intractable problem of optimally accounting for food depletion into a sequence of simpler, solvable problems for each stage.

To formalize this approach, we first divide the planning horizon into *N* “stages” of fixed length *S* = *T* /*N* denoted by [*T*_0_, *T*_1_], [*T*_1_, *T*_2_], …, [*T*_*N*−1_, *T*_*N*_], where *T*_0_ = 0, *T*_1_ = *S*, and *T*_*N*_ = *T*. For simplicity of presentation, we assume that the food density function *ψ*(**x**) does not have any explicit time dependence, but the model supports time-dependent food densities as well. For stage *n ∈* {0, …, *N* − 1}, mode *m ∈* {1, 2}, and *s ∈* [0, *S*] we define the value function 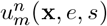 which represents the optimal reward-to-go until the end of stage *n* from the current state and mode. To compute these value functions we alternate between solving a version of equations (8) and (9) over the interval *s ∈* [0, *S*] and simulating a realization of the forager’s optimal trajectory over a single stage, updating the food availability map based on that trajectory. To initialize, we solve for 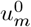 using the initial food density *ψ*(**x**) to compute *F* ^0^(**x**), the food availability during the first stage. Then, we trace a realization of the trajectory **y**(*t*) for *t ∈* [*T*_0_, *T*_1_] using the optimal feedback policy prescribed by 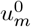.

When tracing the trajectory of the forager, we model its food collection *accounting for food depletion*. An animal truly collecting food according to our model should deplete the available food according

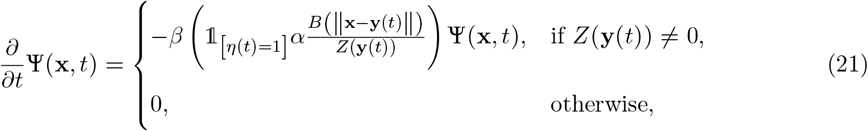

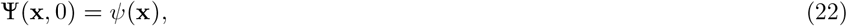

where Ψ(**x**, *t*) is the now time-dependent depleting food density and

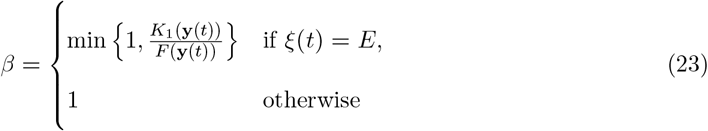

captures the fact that the animal can only eat a limited amount when it is at maximum energy. For a known trajectory **y**(*t*) and a fixed **x**, finding the resulting Ψ(**x**, *t*) is simply a matter of solving a linear ODE. On a grid, the decreasing food availability at all relevant gridpoints **x**_*i,j*_ can be updated more efficiently simultaneously with tracing the trajectory **y**(*t*). It is worth noting, that this trajectory is actually random since the forager’s plans often change due to random encounters with predators. This implies that the trajectory-dependent decreasing food availability map Ψ(**x**, *t*) becomes random as well. Once we have solved for 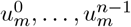 and Ψ(**x**, *t*) for = *t ∈* [*T*_0_, *T*_*n*_], we can compute 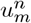 in the same manner by setting *ψ*^*n*^(**x**) = Ψ(**x**, *T*_*n*_), using *ψ*^*n*^(**x**) to compute *F*^*n*^(**x**), and solving equations (8) and (9) on [*T*_*n*_, *T*_*n*+1_]. This assumes that we have chosen a terminal reward for each stage, which we will take to be the expected utility of the forager’s energy at the end of the stage.

We illustrate our multistage approach on a simple environment with two food-rich areas and a safe refuge (Figure 10a, again similar in spirit to a discrete two-patch model). While the food availability and distance to the refuge are identical in both “patches,” we think of the bottom patch as having “cover” that reduces the likelihood of the forager being killed there (Figure 10). We assume that the animal travels with constant speed *f*_*m*_(**x**) = 5, expends energy at a constant rate *K*_*m*_(**x**) = 0.8, and faces constant spotting rate *µ*_*s*_(**x**) = 3 outside of the refuge. Inside of the refuge, the forager cannot be spotted or killed (*µ*_*s*_(**x**) = *µ*_*k*_(**x**) = 0) and the predator is more likely to give up on the chase (*µ*_*g*_(**x**) increased by a factor of 10). Since the two patches are otherwise identical, a rational forager without knowledge of food depletion would only exploit the bottom patch (assuming it starts within the refuge) and would thus do poorly if significant food depletion actually occurs. However, a depletion-aware forager may sometimes exploit both regions, though still favoring the lower.

**Figure 10:**
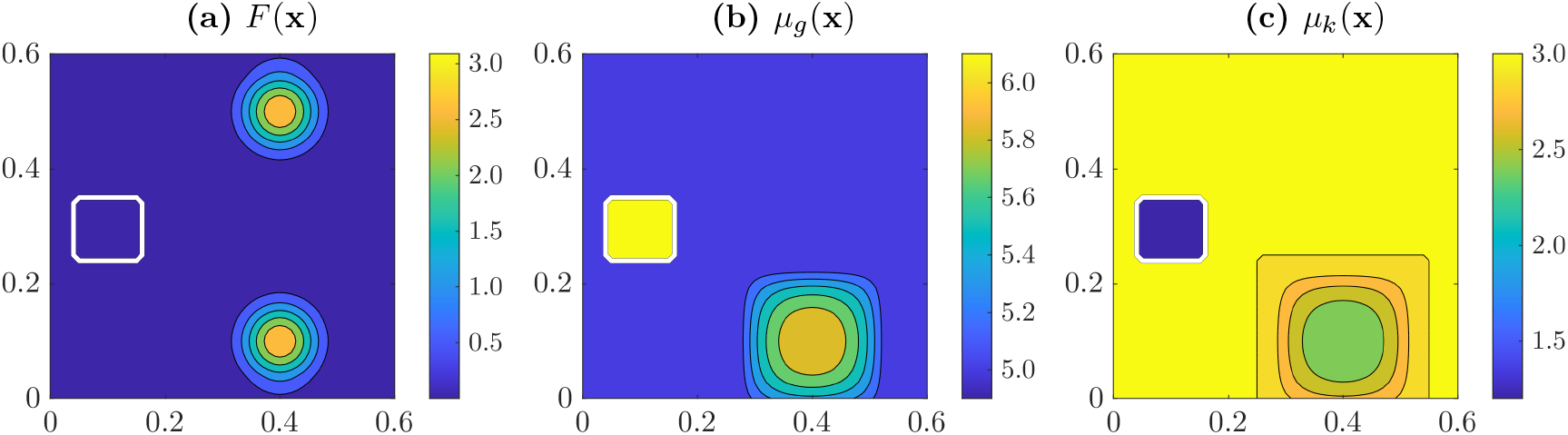
A “two-patch” environment where one patch is strictly better than the other. Two regions of equal food availability are equidistant from the refuge (subfigure **(a)**). The lower patch is assumed to have cover, which increases the chance of escaping (subfigure **(b)**) and decreases the chance of being killed (subfigure **(c)**) when there is a predator. A rational forager that starts in the refuge and does not deplete food will only exploit the lower patch, but our “multistage model” correctly predicts that it can be optimal to exploit both patches when depletion occurs (see Figures 11 - 13). Exact expressions for each quantity are provided in Section E.

Figure 11 shows three realized trajectories under the multistage model for a risk-neutral forager. If not spotted by a predator (top row), the forager spends the first three stages exploiting the safer food patch. At the beginning of the fourth stage, the patch has been depleted to the extent that it is optimal to travel to the unexploited top patch, though taking a detour to remain closer to the refuge. If the forager is instead spotted while traveling between patches (middle row), it escapes towards the refuge to avoid the predator. When the forager is later spotted near the end of stage four, it misjudges the cumulative danger since it does not plan beyond the end of that stage and the remaining time is insufficient to reach the burrow anyway. As a result, it temporarily lingers in that patch (despite the predator’s presence) and only returns to the refuge once the next stage begins. This behavior is an artefact of the modeling process and not biologically motivated. The bottom row of Figure 11 highlights a possible impact of an early predator spotting: when spotted at the end of stage two, the forager escapes to the refuge in stage three, and then takes advantage of its central position to travel to the unexploited upper patch. As a result, it arrives in the risky patch earlier, and spends more time there than in the safe patch over the total time horizon.

**Figure 11:**
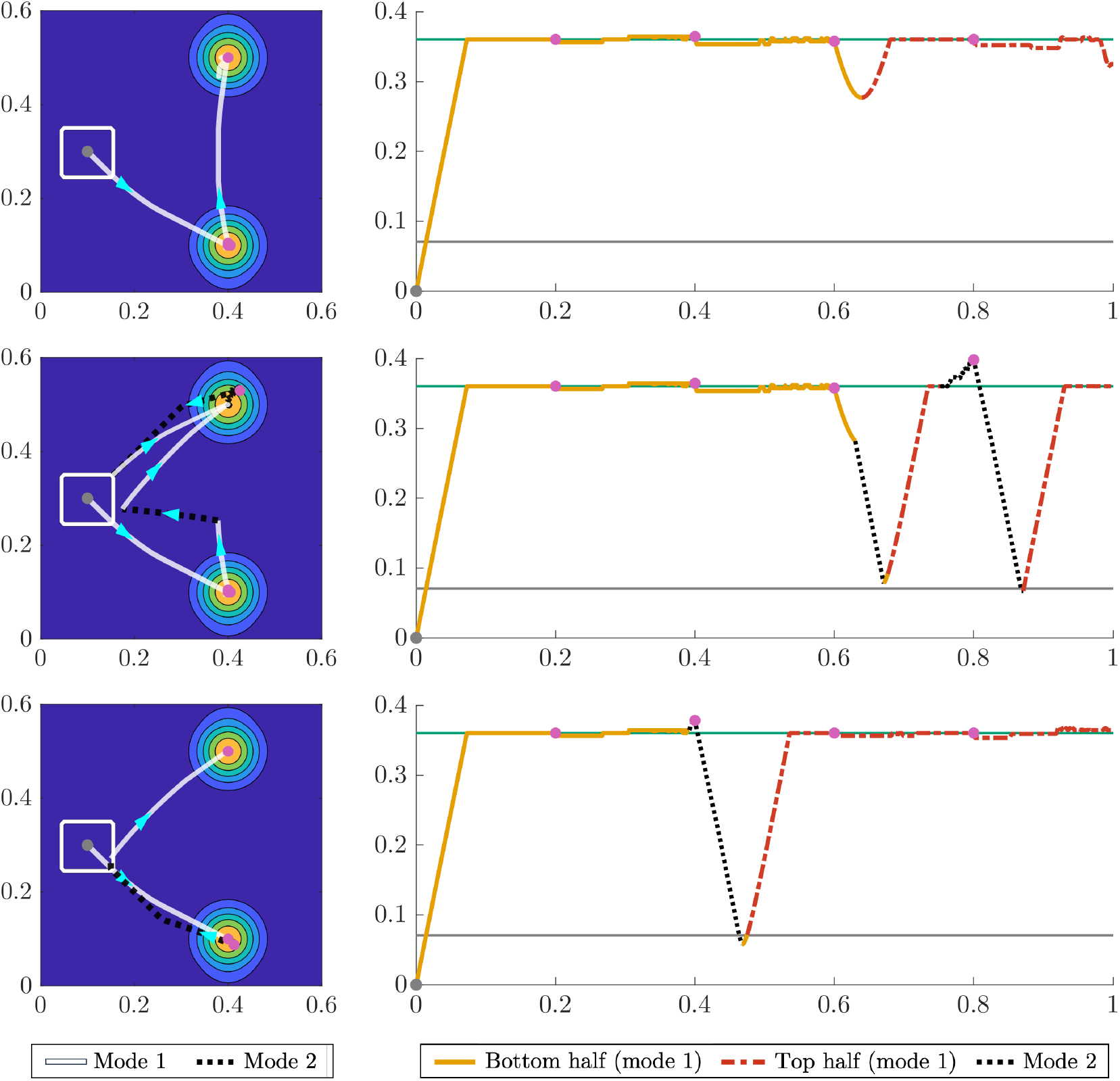
A myopic forager that periodically becomes aware of food depletion exploits both food patches (Risk-neutral). Three realized optimal trajectories according to the “multi-stage model” with five stages using a risk-neutral (linear) utility function. The forager begins in the center of the refuge (grey dot) and the end of each stage is marked with a magenta dot. The first column shows optimal trajectories in the forager’s 2D environment, with blue arrows indicating the direction of travel. In the second column, we plot the forager’s distance from the starting location over time for the same realized trajectories. The green line represents the distance to the center of each food patch and the grey line is the distance to the *corners* of the refuge. Across all realizations, the forager preferentially exploits the safer food patch, but eventually switches to the top patch as it becomes aware of the impact of food depletion. If not spotted by a predator, the forager travels directly from the first patch to the second (top row). However, if spotted by a predator it may return to the refuge (middle and bottom rows), and in some cases switch to the top patch earlier due to such a detour (bottom row). “Wobbling” in the trajectories is caused by small variations in food availability due to depletion.

Figures 12 and 13 present similar stories but for risk-averse and sigmoid planners. In the absence of predators, the behavior of the risk-averse planner is similar to that of the risk-neutral planner, though it takes a longer detour between the patches to remain close to the refuge (Figure 12, top row). When not spotted by a predator, the sigmoid planner only exploits the safer patch before returning to the refuge to wait out the end of the planning horizon. However, it if travels to the refuge to escape a predator or loses foraging time due to a chase it may exploit the upper patch as well (Figure 13, middle and bottom rows). In this setting, the multistage model can capture more nuanced behavior than a model without food depletion, despite the limitations associated with stage length and terminal condition.

**Figure 12:**
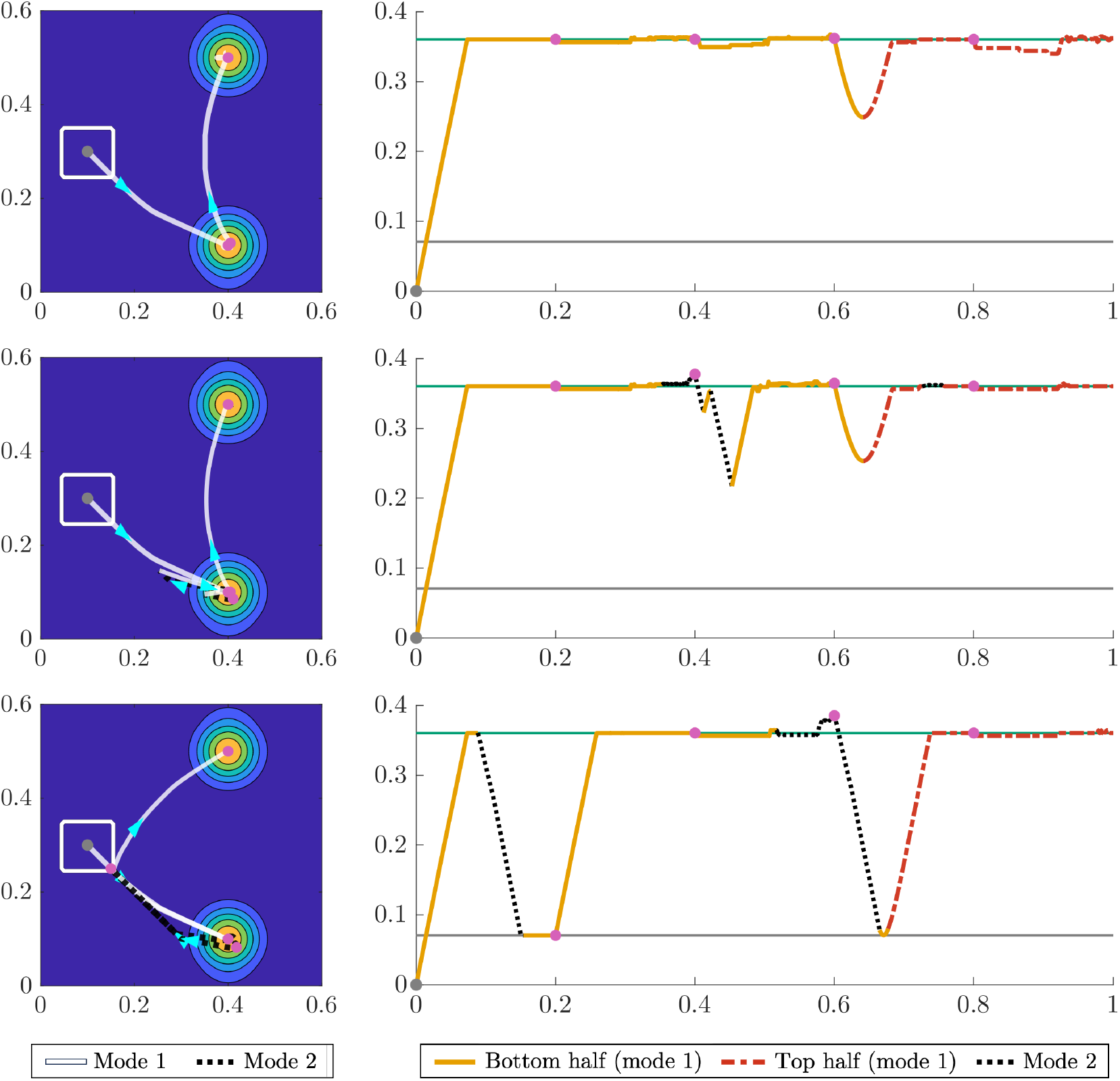
A myopic forager that periodically becomes aware of food depletion exploits both food patches (Risk-averse). Formatted as in Figure 11. In the absence of predators, the behavior of the risk-averse planner is similar to that of the risk-neutral planner, though it takes a longer detour to remain close to the refuge on the way to the second food patch (top row). If the forager is spotted by a predator but successfully escapes before reaching the refuge, it may return to its original foraging patch (middle row). In the third row, the forager escapes to the refuge twice, the first chase is early enough that it is optimal to return to the lower patch (bottom row). After the second chase, the forager switches to the upper patch due to the effects of food depletion.

**Figure 13:**
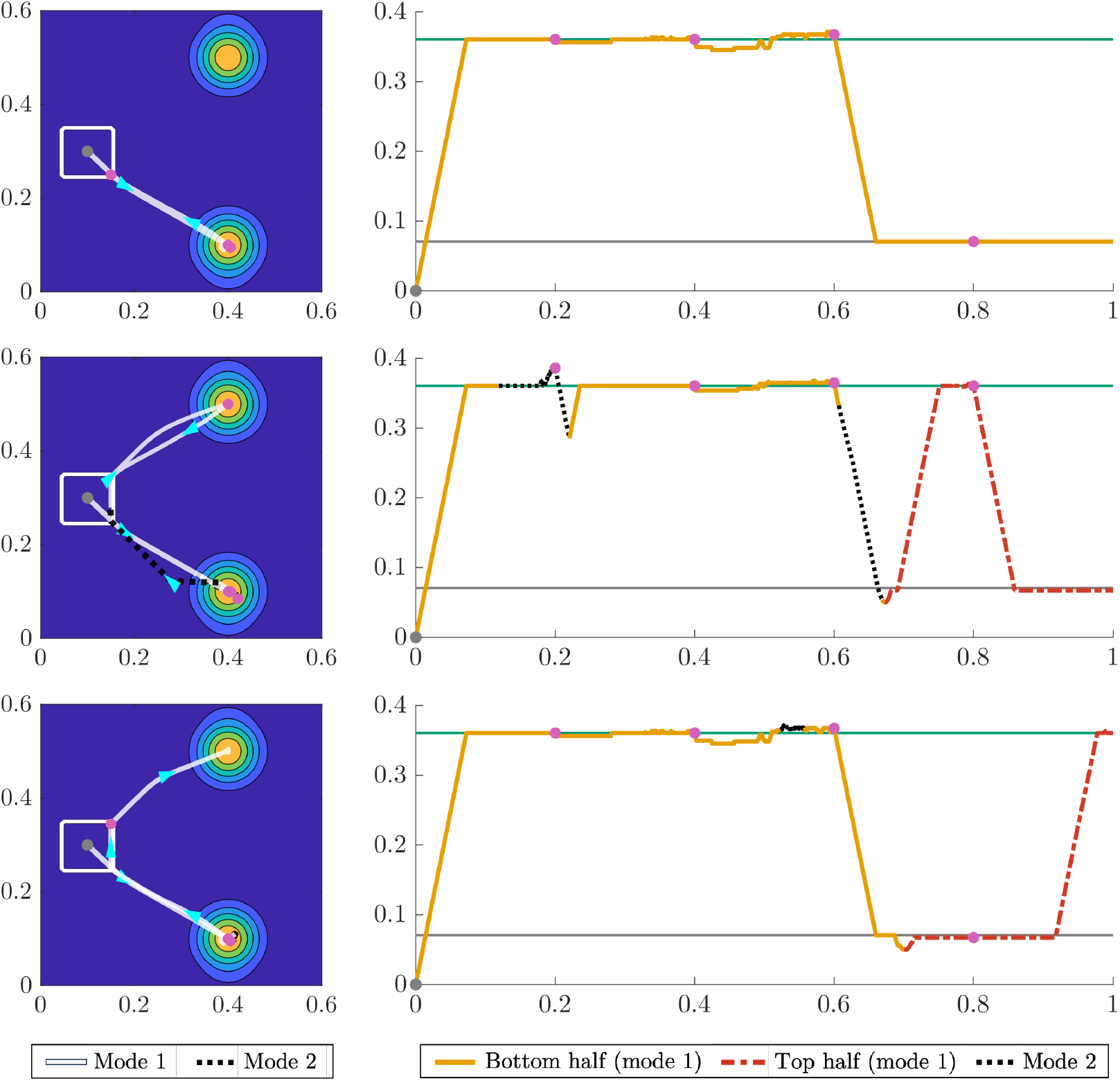
A myopic forager that periodically becomes aware of food depletion exploits both food patches (Sigmoid). Formatted as in Figure 11. The sigmoid planner is the most cautious of the foragers we model. In the absence of predators, it only exploits the safer patch (top row) before returning to the refuge at the beginning of the fourth stage, a clear distinction from the other two planners. However, the arrival of a predator can cause the forager to return to the refuge earlier than planned, and from there is may be optimal to switch to the upper patch (middle row). In the bottom row, a forager that has only a brief encounter with a predator later chooses to travel to the upper patch to make up for lost foraging time.

A key strength of the multistage model is that it incorporates the forager’s feeding history in a computationally feasible manner. Indeed, the cost of solving the dynamic programming problem is the same as for the original model. However, because these solutions incorporate information specific to a single trajectory, they cannot be used to efficiently simulate trajectories for many starting locations or for many different random interactions with predators. Furthermore, the multistage model introduces additional modeling choices: the terminal condition and length for each stage. The forager’s expected utility at the end of the stage is a natural terminal condition, as we have already interpreted it as a measure of future reproductive success. However, it does not incorporate any information about the forager’s future prospects in the short term, and may lead to unrealistic behavior, e.g., a forager that stays in place and starves because it cannot reach a food source within a single stage (even if it could reach such food over multiple stages of travel).

A careful choice of the stage length can alleviate this issue, but is often far from obvious. Even when each stage is long enough for the forager to reach food, travel will take up proportionally more time for short stages, exaggerating the opportunity cost associated with traveling. Similarly, the forager will underestimate the risk of being killed by a predator for shorter stages or near the end of a stage, because it is less likely to be spotted and killed before the stage ends (e.g., stage four in the middle row of Figure 11). However, overly long stages inherit the drawback of the original single-stage model, that a forager might remain at a depleted food source when it is better to move on. Thus, stages must be long enough for the forager to reasonably estimate trade-offs between travel, foraging, and predation, but short enough that it receives timely information about depleting food. Despite these limitations, we believe that this approach is an important step in developing approximate models that can better represent situations where foragers impact their environment.

## D Numerical Methods

### D.1 HJB PDE Discretization

We solve equations (8) and (9) backwards in time using an explicit finite-difference scheme. We denote the grid spacing using Δ*x* = Δ*y* = *h*, Δ*e*, and Δ*t*. Further, we denote a point in our discretized (*x, y, e, t*) space as (*x*_*i*_, *y*_*j*_, *e*_*k*_, *t*_*l*_) where *x*_*i*_ = *ih, y*_*j*_ = *jh, e*_*k*_ = *k*Δ*e*, and *t*_*l*_ = *l*Δ*t*. While we use Δ*x* = Δ*y* for simplicity of presentation, it is straightforward to generalize the scheme to Δ*x*≠ Δ*y* as well.

Let *U*_1_ and *U*_2_ represent the solutions to the discretized system of equations, such that

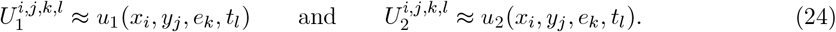

We can then similarly write 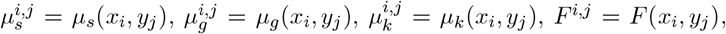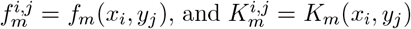.

In addition to solving backwards in time, we use an *upwind* discretization in space and energy. First, we can define the following one-sided difference operators:

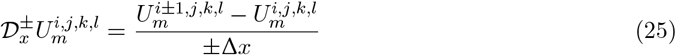

with respect to *x*, where operators for *y, e*, and *t* can be defined in the same way. We can use these one-sided difference operators to construct upwind difference operators for the *x, y*, and *e* coordinates to ensure that the optimal trajectory is always straddled by the computational stencil. Starting with *x* and *y*, the upwind operators are defined as

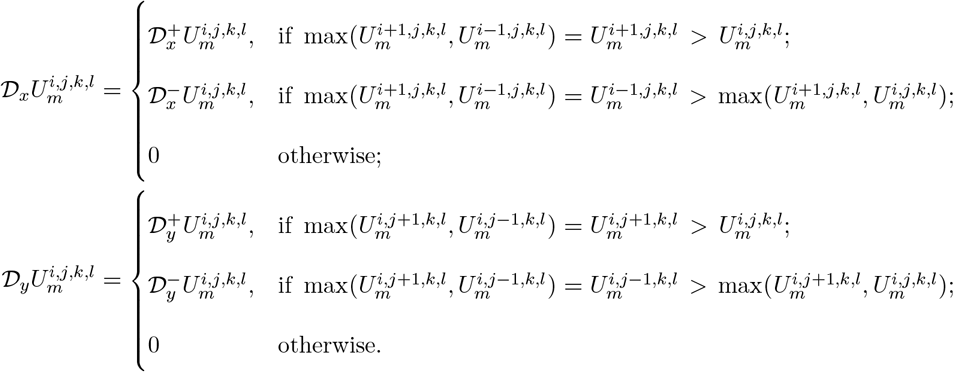

For the energy dimension, the upwind operator can be defined a priori as long as the mode is known. If *m* = 1 the animal collects and expends energy, so the upwind direction depends on the sign of 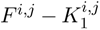. Since both functions are known, we can define the following upwind operator:

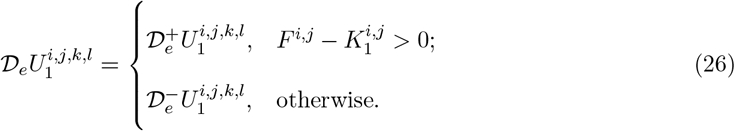

If *m* = 2 then the animal cannot collect food and the energy level must be non-increasing so we simply have

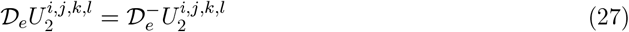

as our upwind approximation.

Putting all of this together, we can write the fully discretized equations as

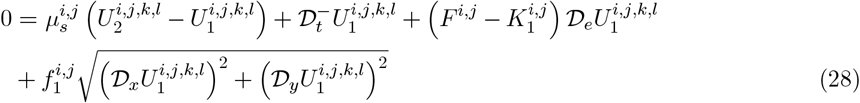

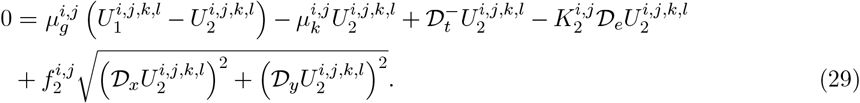

We use 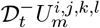 as the derivative approximation in the time dimension in order to obtain an explicit update for 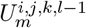 in terms of quantities in the *l*th slice of the value function. Rearranging equations (28) and (29) we get the following update rule:

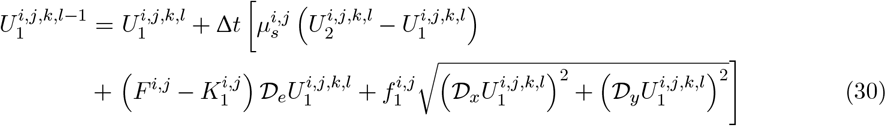

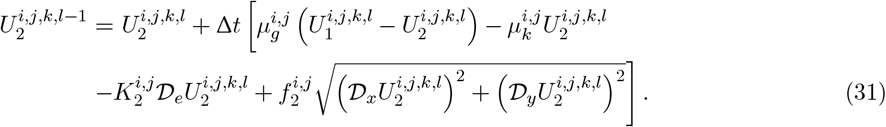

### D.2 CFL Conditions

Since we are using an explicit discretization in time, the stability of the numerical scheme is governed by a Courant-Friedrichs-Lewy (CFL) condition. This condition is based on the dynamics of the problem, and since each mode has its own dynamics, each will also have its own CFL condition. For simplicity, we use the same time step for both modes in our numerical scheme, so our choice of Δ*t* is based on the more restrictive of the two conditions. We now derive the CFL conditions for modes 1 and 2. **Mode 1:** The analytic domain of dependence for a gridpoint (*x*_*i*_, *y*_*j*_, *e*_*k*_, *t*_*l*_) at the slice *t* + Δ*t* is the circle in (*x, y, e*)-space centered at 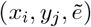 with radius *R*, where 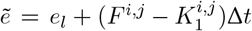 and 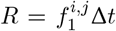. For the CFL condition to be satisfied, the above circle must lie within the eight simplexes generated by (*x*_*i*_, *y*_*j*_, *e*_*k*_) and its 6 neighboring gridpoints. To simplify the notation, we will for the moment shift our coordinates so that (*x*_*i*_, *y*_*j*_, *e*_*k*_) = (0, 0, 0) and omit the superscripts. Without loss of generality, we assume that (*F* − *K*_1_) *>* 0 and focus on the simplex in the first octant. The relevant part of the boundary of that simplex is given by 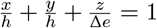, where *h* = Δ*x* = Δ*y*. We now compute the maximum radius of a circle centered at 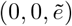 parallel to the *xy*-plane lying within the simplex. In the intersection of the plane 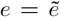 with the simplex boundary in the first octant, 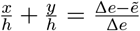. So the intersection of this plane with the simplex is a right isosceles triangle with leg lengths 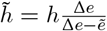. Thus, the distance from (0, 0) to the hypotenuse gives us the maximum allowable radius 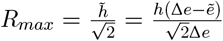. Rearranging *R ≤ R*_*max*_ and maximizing over all gridpoints, we find

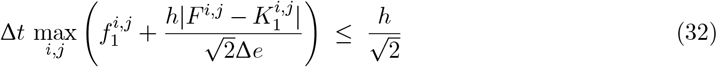

(In a more general case, where *K*_1_, *F*, or *f*_1_ could depend on the current energy state or time, we would also maximize over *k* and *l*.)

#### Mode 2

This is the same as mode 1, except the (absolute value of) rate of change of energy is | − *K*_2_| = *K*_2_, and we use *f*_2_ instead of *f*_1_ for the speed in physical space. Thus the CFL condition becomes

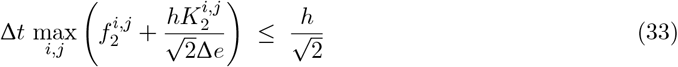

Combining mode 1 + mode 2, we find the combined CFL condition

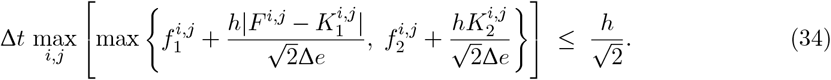

### D.3 Path Tracing and Food Depletion

To trace the optimal trajectories, we simulate the position and energy dynamics using a forward Euler scheme with time step *τ* (which may differ from Δ*t* used to solve the PDEs). We use a grid search over directions of motion to determine the optimal direction, *θ*. Slightly abusing notation, if (*x*_*l*_, *y*_*l*_, *e*_*l*_, *t*_*l*_ = *lτ*) is the approximate state of the forager after *l* steps of size *τ*, for motion in the direction (cos *θ*, sin *θ*), the next position is approximated as

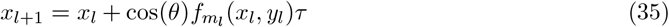

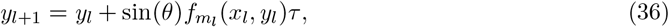

and the actual optimal direction of motion is selected as

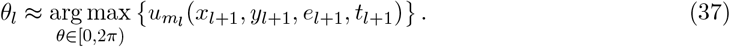

For energy, without food depleting, the update rule is given by

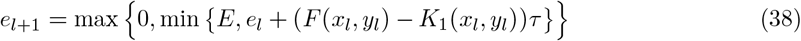

in mode 1 and

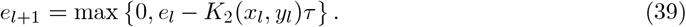

in mode 2. We linearly interpolate all functions when evaluating them between gridpoints, except for *F* (**x**), which we compute via numerical quadrature for each location.

When considering food depletion, the update for the energy is more complicated. To compute *e*_*l*+1_ in mode 1, we deplete the food in the environment according the animal’s position (*x*_*l*_, *y*_*l*_). To deplete food at a given gridpoint **x** = (*x, y*), we consider two cases. First assume that the animal cannot reach *E* (in time Δ*t*) from its current energy level. Because of the CFL condition governing Δ*t*, this is equivalent to the condition *E* − *e*_*l*_ *>* Δ*e*. In this case, we approximate the solution to equation (21) on the small interval [*t*_*l*_, *t*_*l*+1_] using a forward Euler step

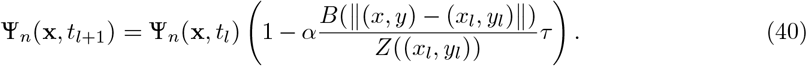

If *E* − *e*_*l*_ *≤* Δ*e*, it is possible that the animal will reach *E* in the next time step, and we instead use the update

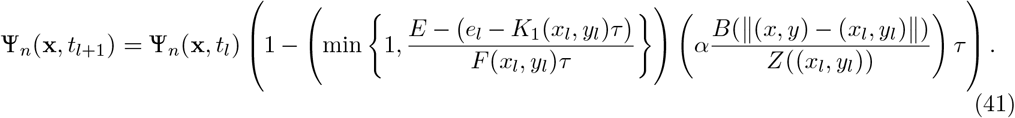

We make use of the fact that the food depleted across the entire environment is equivalent to the food accumulated by the animal, and set

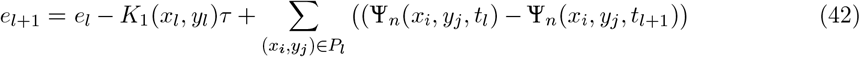

where *P*_*l*_ is the set of grid points for which Ψ_*n*_(*x*_*i*_, *y*_*j*_, *t*_*l*_)≠ Ψ_*n*_(*x*_*l*_, *y*_*l*_, *t*_*l*+1_). In mode 2 the animal does not eat so there is no food depletion and we can use the update from equation (39).

Since we use a forward Euler step to simulate the ODE for Ψ_*n*_, this imposes an additional restriction on the time-step to ensure stability. We must choose *τ* such that

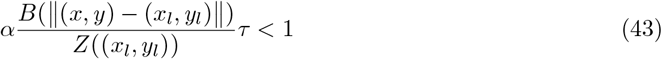

In practice, we assume that *B* is continuous and nonincreasing on [0, *r*] and *τ* takes the form *τ* = ^Δ*t*^. We can then substitute the stricter condition

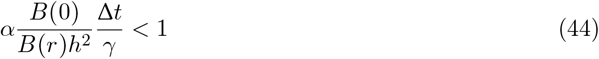

and choose *γ* such that

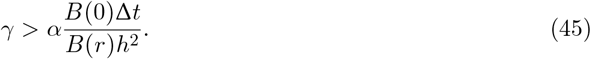

## E Formulas for Numerical Experiments

### E.1 Balancing Risk and Reward Environment

In this example, we compute the food availability based on food density

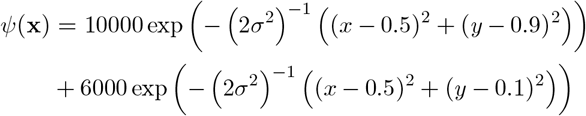

with *σ* = 0.05. We assume that the animal has food radius *r* = 0.01, harvest rate *α* = 0.001, and that the food kernel *B*(*d*) is the characteristic function of *D*_*r*_(0). The animal’s initial position is (*x*_0_, *y*_0_) = (0.41, 0.5). For a fixed risk premium *ρ*, the spotting rate is computed as

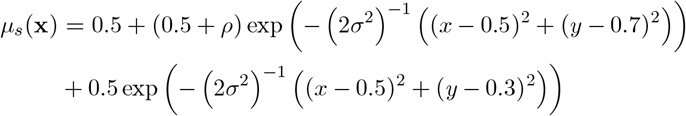

for *σ* = 0.1.

### E.2 Real-world Inspired Environment

In order to compute the food availability from our estimated food density, we require two additional parameters: a radius *r* that describes the local area in which the monkeys gather food, and a harvest rate *α* that describes how efficiently monkeys can collect food in their environment (see equation (12)). For this example, we use *r* = 0.05 km and *α* = 10^−6^ h^−1^.

### E.3 Multistage Model Environment

We compute the food availability based on food density

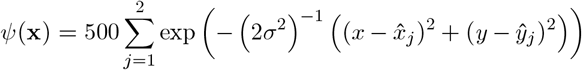

for 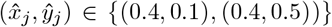 and *σ* = 0.035. We further assume that the animal has food radius *r* = 0.05, harvest rate *α* = 0.01, and that the food kernel *B*(*d*) is the characteristic function of *D*_*r*_(0). For all experiments, we set the initial energy to be *e*_0_ = 1.0 and the initial position to (*x*_0_, *y*_0_) = (0.1, 0.3). The give up rate and kill rate are computed as follows:

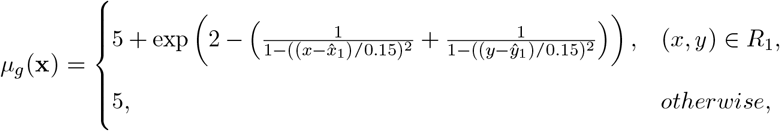

and

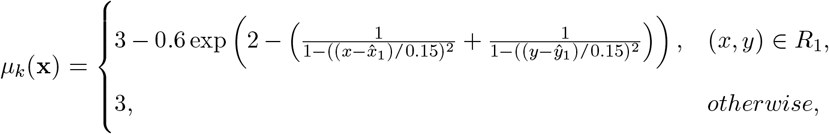

where *R*_1_ is the square of width 0.15 centered at (0.4, 0.1).

## Footnotes

1 These are *isotropic* dynamics, which can represent how speed depends on local vegetation density or other location-based terrain characteristics. Anisotropic dynamics (e.g., to model the dependence of speed on the terrain slope in the chosen direction) can be handled similarly, yielding more general Hamilton-Jacobi-Bellman PDEs. While the general computational approach remains the same, it would require more data to calibrate the model and we choose this simpler setting to highlight the main ideas.

2 This comparative approach is, unfortunately, uncommon in the literature. But we note that [Newman, 1991] also examines the impact of the shape of a utility function on foraging behavior, though for a discrete patch-based model with depletable resources.

3 More formally, we represent each region of elevated risk using a Gaussian function, and define the “risk premium” *ρ* to be the difference in their peak values. See Supplementary Materials E for detailed expressions.

4 Here, the preferred patch is the patch that the animal initially plans to visit before any possible predator encounters have occurred. If the forager is spotted by a predator en route, it may return to the refuge before reaching its preferred patch and may visit a different patch on a subsequent excursion.

5 Recall that food density is used to compute food availability *F* (x). See Supplementary Materials Section A for the details of this calculation.

